# A microfluidic approach to explore mesoderm tissue dynamics and its natural variability

**DOI:** 10.64898/2026.04.22.720163

**Authors:** Guillaume Desgarceaux, Majid Layachi, Christine Fagotto-Kaufmann, Laura Casanellas, François Fagotto

## Abstract

Vertebrate gastrulating mesoderm is a prototypic example of a mesenchymal-like tissue undergoing extensive remodelling. While the tissue may be globally represented as a viscoelastic material, the actual biological material is intrinsically complex. To get to a real understanding of its properties, one needs to move to the mesoscale, linking cellular properties to collective phenomena. Vertebrate embryos also display a remarkable variability in mechanical properties, despite which they robustly complete gastrulation. This study attempts to explore these aspects by dissecting Xenopus mesoderm cell behaviour in a minimal system, using aspiration through a microfluidic system to impose controlled stress to a mesoderm aggregate. We show that beyond estimating global rheology at the tissue scale, it is possible to infer a wealth of information based on cell morphology and dynamics. Our data are consistent with collective behaviour being mostly dictated by the balance between the capacity of cells to stretch and the resistance to cell-cell contacts, which limits cell-cell intercalation and thus tissue remodelling. Importantly, tissues are not only able to transmit stress over a distance, they also clearly react to it through actively reinforcing cell-cell mechanical coupling. This adaptative property is found through a broad range of tissue stiffness, and adhesion strength appears to scale with the elastic modulus, suggesting that cell stiffness may ultimately be the key parameter setting mesoderm rheology and accounting for the large differences observed between embryo batches.

## Introduction

During development, morphogenetic events position and shape embryonic structures which will later give rise to the tissues and organs of the adult. Internalization of the mesoderm during Xenopus gastrulation is an archetype of massive collective migration of a multilayered, highly coherent tissue. This process, called involution, is mechanically challenging as the cell group has to make its way through the compact embryo. Several types of forces are involved (Huang and Winklbauer, 2018; Keller and Winklbauer, 1992): Cells contacting directly the substrate exert a pulling force as they advance. Mesoderm cells mostly crawl directly on the surface of ectoderm cells, with a minor contribution of a sparse extracellular matrix. Cells within the migrating mass also crawl using their neighbors as substrates, which is an effective power that allows the migrating mass to self-propel. As cells migrate relative to each other, they must eventually remodel their cell-cell contact to exchange neighbours, a phenomenon called cell intercalation (Walck-Shannon and Hardin, 2014). Cells not only exert forces, but also experience external forces, including pulling forces transmitted by fellow cells - those moving ahead and those trailing behind, as well as compressive forces generated as the cell mass squeezes between other tissues. Numerous feedback reactions occur at all levels, as cells sense, react and adapt to intrinsic and extrinsic forces, thus forming a complex active material. While in the embryo, mesoderm involution is partly guided by attractive cues (Damm and Winklbauer, 2011; Nagel and Winklbauer, 2018), migration is intrinsic to mesoderm cells and is maintained in isolated explanted tissues.

In contrast to classical examples of stereotyped and beautifully synchronized morphogenetic processes extensively studied in invertebrates, in particular in Drosophila, a striking and puzzling aspect of vertebrate gastrulation, including mesoderm involution, is the heterogeneity of the tissues and the huge variability between embryo batches, both at the tissue and cell scales, reflected in parameters such as stiffness, adhesiveness, or migration speed (Blanchard et al., 2019; Canty et al., 2017; Damm and Winklbauer, 2011; David et al., 2014; Huang and Winklbauer, 2022; Kashkooli et al., 2021; Luu et al., 2008; Luu et al., 2011; Rozema et al., 2025; von Dassow and Davidson, 2009; von Dassow et al., 2010). How embryos nevertheless manage to perfectly complete gastrulation is a fascinating question that has yet to be understood.

Dynamic tissue can be modelled based on global physical properties, and generally appear to behave as a viscoelastic material (Gonzalez-Rodriguez et al., 2012; Marmottant et al., 2009; Phillips et al., 1977), a description that holds for Xenopus mesoderm (David et al., 2014; Luu et al., 2011; von Dassow et al., 2010). This coarse grain phenomenological description of a tissue must be related to the behaviour of its cell components. Indeed, cells constitute the basic autonomous units, and tissue rheology is the emergent result of the underlying properties of the cells and of their interactions. Typically, elasticity of the tissue is viewed as resulting from the capacity of the cells to reversibly deform, provided that cell-cell adhesive connections resist to the load and maintain the integrity of the tissue. If adhesion fails, cell contacts may break, making the tissue more “liquid”. However, cells have the capacity of maintaining tissue integrity through dynamic contact remodelling, releasing existing contacts and forming new ones, which is the essence of the process of intercalation. In this case, the tissue will irreversibly deform, while remaining coherent. The capacity of cell-cell contacts to resist remodelling, thus impeding contact exchanges, represents the viscous component of this cellular material (Marmottant et al., 2009). Note that, conversely, active cell processes that stimulate intercalation can in principle decrease apparent viscosity. In this study we have been interested to set an assay that could bridge tissue scale properties of embryonic tissues with the properties of its cell constituents, specifically in the context of a multi-layered mesenchymal tissue, using as model the Xenopus anterior or prechordal mesoderm. We have used a microfluidic approach to study small mesoderm aggregates under stress and confinement, coupling biophysical estimation of tissue rheology with live imaging of the dynamics of individual cells. From these exploratory attempts, we propose that despite the daunting complexity of collective behaviour, it is possible to get a good grasp at the process by interpreting changes in cell morphology based on simple principles of tension and adhesion.

## Results

### Experimental setup

The microfluidic device was adapted from a previous design used to study synthetic adhesive lipid vesicles (Layachi et al., 2025), and was similar to the one developed by Graner and colleagues to study aggregates of mammalian culture cells (Tlili et al., 2022). Figure 1A presents the detailed design of the device. The setting comprises a loading inlet followed by a 250μm-wide pre-chamber leading through a 45° funnel-shaped constriction to a narrow (100μm) channel. The design was symmetrical (Fig.1A), but for here we only monitored entry in the channel. The funnel was included both to avoid potential re-circulation zones at the corners, and to impose a progressive constraint mimicking what tissues may experience, in particular the mesoderm entering through the blastopore lip in the intact embryo. In straight channels the applied pressure difference creates a purely shear flow (with the highest shear rate being localised next to the channel walls). In the contraction geometry, instead, the flow is heterogeneous and comprises both shear and extensional contributions. The rheological parameters that we will obtain hereafter are thus apparent quantities, as they arise from a combined shear and extensional solicitation rather than a purely single-mode deformation. Mesoderm cells where dissociated from explanted mesoderm, loaded, gently aspirated in the pre-chamber and left to reaggregate. Our well-established dissociation conditions preserve very well cell-cell adhesion, cells being able to re-establish adhesive contacts within few minutes (Kashkooli et al., 2021; Rohani et al., 2014; Rozema et al., 2025). Once they had formed a compact mass (~30min), they were aspirated through a funnel-shaped constriction and the narrow channel. Importantly, low aspiration pressures were used, which started in the range of 5 to 25Pa, and were progressively increased. This strategy contrasts with previous pipette aspiration experiments by others and ourselves, where a high pressure was immediately applied (Kashkooli et al., 2021; Petridou et al., 2019). Such harsher conditions caused a quasi-instantaneous phase of tissue deformation, likely due to abrupt disruption of cell-cell contacts. For comparison, the time of spontaneous contact remodelling in the Xenopus mesoderm is typically in the minute range (Rohani et al., 2014; Rozema et al., 2025). The chamber had an even height of 100μm, corresponding to a tissue thickness to 2-3 cells. The aim was to limit vertical intercalations, such that the tissue behaved as an almost 2D system. Expression of membrane targeted GFP allowed to directly monitor the morphology and behaviour of the cells within the aspirated tissue. The signal varied in intensity and was often mosaic, but was sufficient to visualize contours of a large number of cells with good precision (Fig.1). Individual tissues will be referred by a number preceded by # and colour-coded throughout the figures (Fig. 1G).

**Figure 1.**
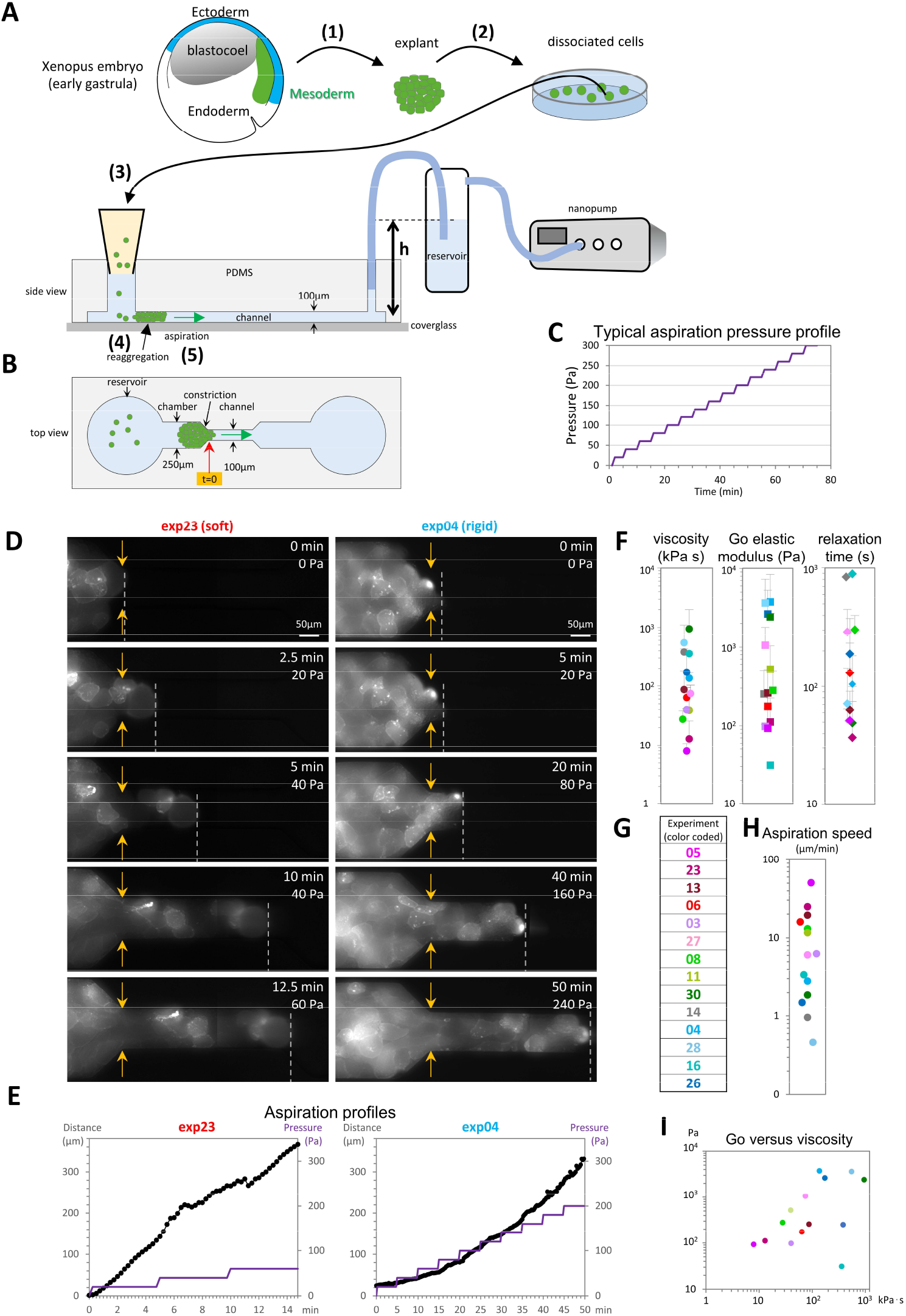
**A-C) Design of the experiment. A)** Preparation of the biological material. Mesoderm tissue was micro-dissected from early gastrula embryos (1), and dissociated into single cells (2). **B)** Side and top view of the microfluidic device symmetrically made of a circular loading pre-chamber where dissociated cells were loaded (3), a 250μm wide chamber where cells were left to reaggregate (4), then aspirated through a funnel-shaped constriction leading to a 100μm wide channel (5). The aggregate was brought just before the entrance of the narrow channel, and default pressure set to 0. Pressure was then set to 20Pa at the start of the time lapse (t=15sec). Images were taken every 15 seconds. **C)** Pressure profile, typically incremented by 20Pa every 5 minutes. **D-E) Examples of aspiration experiments for a soft and a rigid tissue explants. D)** Selected frames from time lapse supplemental Movies 1 and 2. Orange arrows point to entrance of narrow channel. Dotted line marks the front of the tissue. **E)** Corresponding aspiration profiles, distance measured from the channel entrance, and pressure profile. **F-I) Estimated rheology values. F)** Viscosity, elastic modulus and relaxation times. Error bars, SD. See supplementary Figure S1 for fitting model, examples of curve fitting and detailed results. **G)** List of experiments with color-code used throughout the figures. **H)** Average speed of the advancing front over the whole movie. **I)** Elastic modulus tends to scale with viscosity.

### Tissue behaviour and rheology

A coherent tissue can be described as an active viscoelastic material. The elastic component is provided by the ability of the cells to reversibly stretch under tension, combined with the supracellular mechanical coupling provided by the cell-cell adhesive contacts. The viscous component can be related to the property of cells to exchange neighbours through intercalation events, moderated by the resistance of cell-cell adhesions. The latter in turn is controlled by the complex association with the cytoskeleton. All these aspects are impacted by active cellular mechanisms, which can lead to deviation from standard models of material rheology, for instance through processes of adaptation or reinforcement.

The classical aspiration method used to characterize rheological parameters involves applying a constant pressure and monitor the advance of the front of the tissue over time. One typically obtains a curve that can be decomposed in an initial fast exponential phase, dominated by the elastic component, followed by a slower linear phase, which reflects the viscous aspect of the material (Guevorkian et al., 2010). We used here a protocol that imposed progressive stepwise increments in pression (20Pa every 5 min in most experiments, figure 1C). The goal was to challenge the tissues with increasing stress, while leaving sufficient time between steps to distinguish the elastic and viscous aspects at the tissue scale. Under these conditions, we observed a continuous advance of the tissue through the channel, allowing to observe cell behaviour, including intercalations, over long periods (Fig.1D,E, Movies 1,2). The average speed of aspiration of the tissues through the channel was in the 1µm/min to 10µm/min range (Fig. 1H). For comparison, free migration of explanted mesoderm cells on fibronectin is in the 2-5µm/min range and advance of the whole mesoderm migration front during gastrulation embryo was estimated to be around 2µm/min (Kashkooli et al., 2021; Luu et al., 2008). Thus, aspiration kinetics in our experiments were physiologically relevant. Curve segments delimited between pressure steps could be fitted by physical modelling to extract apparent viscosity and elasticity of the cell aggregate, specifically the viscous modulus, the elastic modulus, and the relaxation time, which sets the transition between the two different flow behaviours. The model, details of the fitting and values for individual experiments are presented in supplemental Fig. S1. Note that only a subset of these segments could be fitted with sufficient confidence, and some steps were fitted with a purely viscous model. Indeed, the tissues showed non-standard behaviour, with phases of stalling and/or retraction (see Fig.1E and Supplemental S2). Such behavior reflects the complex properties of these active cell populations. Some of the features that account for these properties will be analyzed below. Thus, estimated parameter values should be taken with caution. Average calculated viscosity modulus, elastic modulus and relaxation for each tissue are given in Fig.1F. The range of values was broad, as predicted from the diversity of behaviours observed during aspiration, and consistent with the documented high variability in cell cortical tension, cell-cell adhesiveness, and global tissue stiffness and cohesiveness between embryo batches (Canty et al., 2017; David et al., 2014; Kashkooli et al., 2021; Luu et al., 2011; von Dassow and Davidson, 2009). The values fell within the range of previous estimates (Supplemental Table 1). While the analysed tissue explants spanned a continuum of values, for simplicity of the presentation of the results, we made two rough categories, which we called soft and rigid tissues, based on the pressure required to aspirate them, their apparent elastic modulus and their viscosity. For the tissues for which we could estimate multiple values at different pressures, both viscosity and the elastic modulus appeared globally independent on the applied pressure (Supplemental Fig. S1). Overall, viscosity and elasticity scaled together, with some exceptions (Fig. 1I). As for the relaxation time, these rough approximations consistently fell in the one-minute range. Note that the steps that were fitted with a purely viscous equation provided no relaxation time (which is equal to zero by definition).

### Cell deformation and elastic properties

Cell deformation under stress provides information about cell elasticity. We thus set to analyse the changes in shape of manually segmented cells over the time of aspiration. Two sets of data were obtained from 5 representative tissues (three soft and two rigid). For the first set, we segmented all cells with discernible contours at five identical time points (Fig.2), which allowed to directly compare the tissue explants under identical time and pressure conditions. Cells contacting the lateral wall of the constriction were excluded from the analysis, as these displayed a peculiar phenotype described below. At each time point, cells were grouped in four categories according to their location relative to the channel entrance (Fig.2B). Shear flow was expected to be dominant in both initial and narrow channels, while elongational flow was present in the funneled constriction, and the tensile stresses exerted to the cells was expected to be maximized close to the channel entrance. Note that the two last regions (entrance and narrow channel) had fewer cells, thus the results were less robust. Segmented cell shapes were fitted to ellipses, from which we extracted the aspect ratio (AR) and the orientation (angle α relative to the longitudinal axis of the channel) (Fig.2C). Figure 2E presents the compiled data from all regions, and Figure 2F the details for each region. Starting with the chamber before constriction, and under the initial condition of low pressure (20Pa), we measured a basal cell elongation (median AR~1.5), consistent with an intrinsic anisotropy also observed for single dissociated cells (e.g. (Kashkooli et al., 2021)). Importantly, cells appeared randomly oriented (median α ~45°). A similar anisotropy was maintained in all regions (Fig.2F), with a weak upward trend at high pressures (Fig.2E). Orientation, expressed by α, aligned with the channel for cells closer to the channel AND with increased pressure (Fig.2E, F). At the channel entrance, α dropped below 20° in all cases. For each tissue, this final alignment occurred only at the upper pressure applied, as low as 60Pa for the soft tissues, up to 300Pa for rigid tissues.

**Figure 2.**
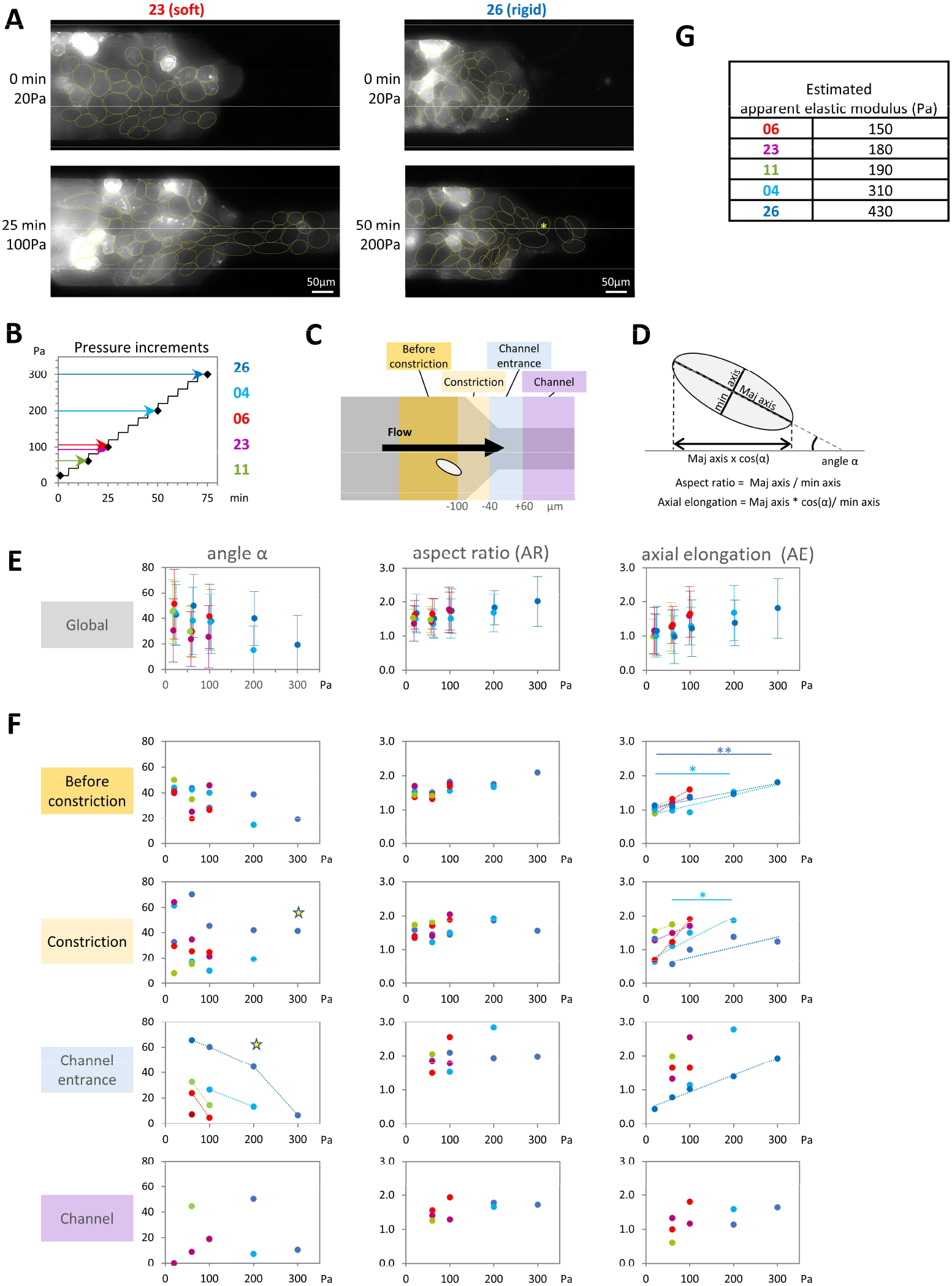
Global analysis of cell shape and deformation. Images from 5 experiments were analysed at frames t=0, 15, 25, 50, and 75min, corresponding to incremented pressures 20, 60, 100, 200 and 300Pa. These time points are indicated by five diamonds along the pressure scale of panel **(B)**, with coloured arrows indicating the highest pressure reached for each explant. The duration of each movie is indicated in D by an arrow. The softest tissues were aspirated at low pressures, thus yielding shorter movies. **A)** Examples of selected frames for a soft #23 and a rigid #27 tissue. Cells were manually segmented and fitted to ellipses. Asterisk: Cell in tissue #27 entering the constriction with a nearly perpendicular ellipse orientation to the flow. **C)** Cells were grouped in categories corresponding to different regions of the microfluid setting. Distance from the channel entrance are indicated below. **D)** Values extracted from the ellipse: aspect ratio, angle α relative to the channel axis, and axial elongation. **E)** Compiled data of all cells until channel entrance included. Error bars, SD. **F)** Median values for α, AR and AE, for each explant, each timepoint/pressure, in the four regions. For a few particularly relevant conditions, the trends are shown by dotted lines, i.e. decreased α at channel entrance, and linear relationship between axial elongation and pressure. Stars highlight the peculiar behaviour for the most rigid explant (27), with cell orientation remaining non-aligned even at high P. Statistical analysis: Two-sided Student’s t-test. **G)** Estimated apparent cell elastic modulus calculated from the slope of axial elongation versus pressure.

We further looked at deformation in the direction of the channel axis, which could be specifically interrogated by calculating the longitudinal component (AR x cos(α)), here defined as “axial elongation” (AE). The shear flow is indeed directed along the axial coordinate in the main and narrow channels. In the constriction, its direction could in principle deviate to a maximum of 45° at the edges, but in the facts, deviations were negligible compared to the variability of cell orientations. Figure 2E shows that AE scaled well with pressure for all tissues (Fig.2E), and this was consistently the case in all locations from before the constriction to the channel entrance (Fig.2F). We could then estimate an apparent elasticity, with a modulus ranging from ~150Pa to 400Pa (Fig.2G). Importantly, the fact that pressure caused axial deformation even before the funnel constriction indicated that stress was efficiently transmitted several cell diameters away from the front of aspiration. On the contrary, once inside the narrow channel, α was less confined, and AR and AE regressed (Fig.2E), indicating that constrains observed at the channel entrance where then loosened. In the thin channel the base flow is purely shear (even though its magnitude exceeds that of the larger channel) with no elongational component, which may explain the observed regression.

This trend was confirmed by single cell tracking: We segmented and tracked single cells through all or most of their path, from before the constriction to the channel (8-14 cells per tissue, for 5 tissues, Fig.3, Movie 3). Figure 3C,D show axial elongation and cell orientation (angle α) for selected individual tracks, with the pressure indicated by a color code. As expected, axial elongation peaked precisely at the channel entrance (Fig.3A,C,E), while α concomitantly dropped to a minimum (Fig.3A,D,F). At the individual cell level, however, a wide range of trajectories were observed before reaching this point: Some cells elongated and oriented far ahead, while others only did so once very close from the entrance. These differences were observed both in soft and rigid tissues. To estimate the impact of aspiration on cell stretching along the channel axis, we plotted axial elongation versus pressure, which gave a strikingly linear relationship (Fig.3G), as expected for a simple elastic system. We used the relative stretching as function of pressure (Fig.3H) to calculate an apparent elastic modulus, which ranged from ~50Pa for the softest to ~300Pa for the stiffest cells (Fig.3I). These values lied within the range of values obtained for the tissues (Fig.2G), and scaled with the estimated elastic modulus of the whole tissue (Fig.1 and Fig.S1).

**Figure 3.**
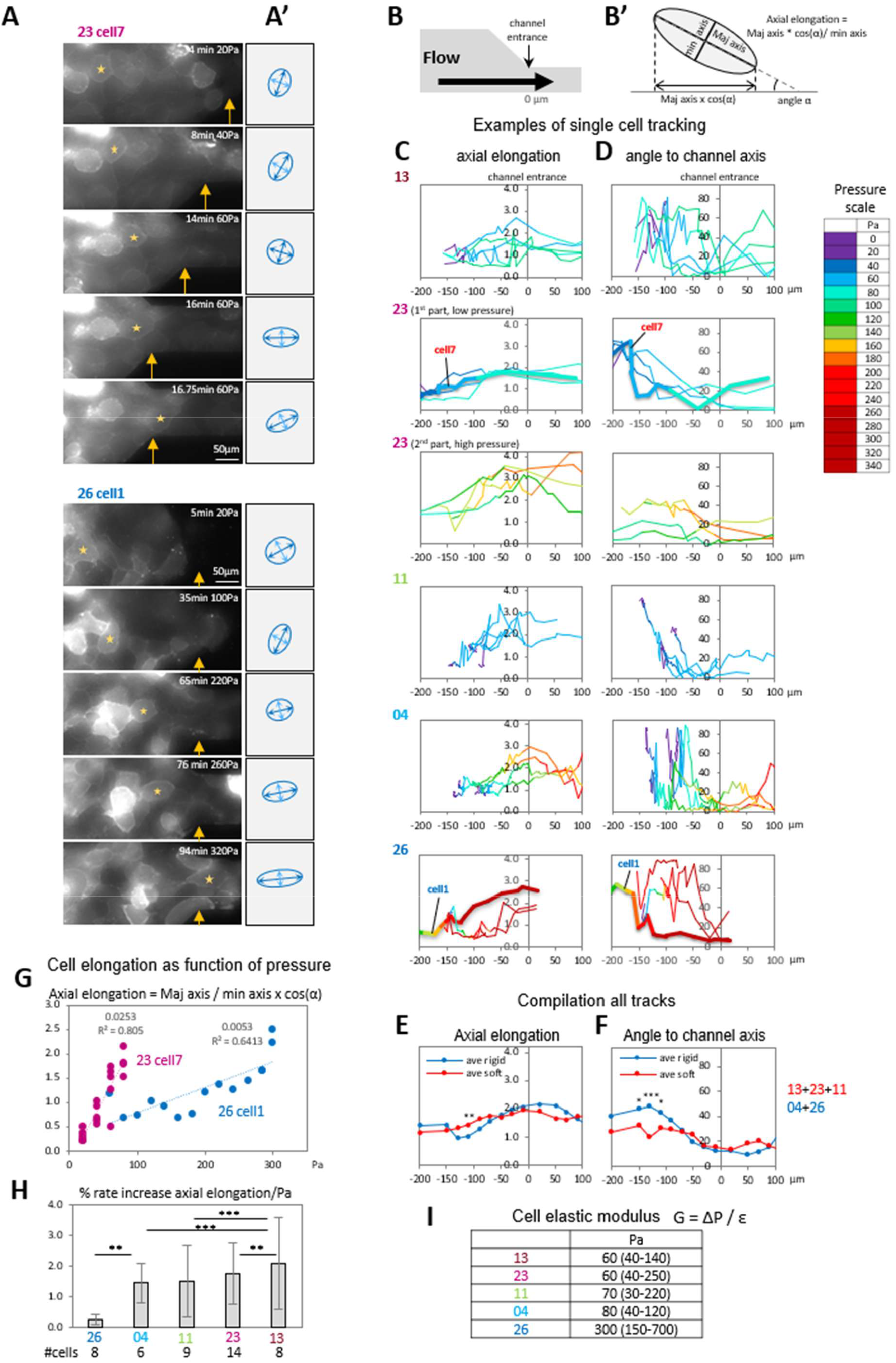
Mapping elongation and reorientation from single cells tracking. A subset of cells from 5 representative experiments were tracked through the time lapse, their outlines segmented and fitted to an ellipse. **A)** Examples of cells in soft (#23) and rigid (#26) tissues, marked by orange star. Detailed field, x position adjusted over time, orange arrows indicate position of channel entrance. See also Movie 3 **A’)** Corresponding ellipses with major and minor axis. **B)** Scheme of half of the chamber, at scale with the x axis of graphs in C and D. **B’)** Scheme of the fitting ellipse with angle α relative to channel axis, and definition of axial elongation. **C-D)** Representative examples of single cell traces, for axial elongation (C) and angle (D). Each trace is color-coded according to the pressure scale. Tissue #23 was constituted of a first soft section followed by a more rigid region. Cells for both regions are shown. Traces corresponding to the cells shown in A) are highlighted by a shadowed thicker line. **E-F)** Compiled data of the three soft and two rigid tissues. In all cases, elongation peaked and angle α reached a minimum very close to the channel entrance. Soft cells tended to show a steadier elongation and become aligned with the channel at a larger distance further ahead of the channel entrance. Statistical analysis: Two-sided Student’s t-test. *,**, ***: respectively p<0.05, <0.01, <0.001. **G-H)** Axial elongation is linearly related to pressure. **G)** Axial elongation at different time points, plotted against pressure, with slope and R^2^ of linear regression. **H)** Average % rate increased (slope/Pa) for the 5 selected tissues. Error bars, SD. Statistical analysis: Non-parametric ANOVA (Kruskal-Wallis Test) followed by Dunn post-hoc test. **I)** Estimated cell elastic modulus, calculated based on H.

### Intercalations and cell adhesive strength

Intercalation is an efficient mechanism to release stress created through the tissue, and one may expect it to vary with the tissue properties, in particular the strength of cell-cell adhesion that opposes to rearrangement (Rozema et al., 2025). In embryos, directional cues orient intercalations, which are otherwise random in tissue explants (Damm and Winklbauer, 2011; Winklbauer and Nagel, 1991). Our microfluidic design limited tissue thickness to 2-3 cell layers, reducing the occurrence of vertical intercalations. This allowed us to focus on the events occurring in the horizontal plane, and analyse them in some detail. We mapped the position of all intercalation events detectable during a time lapse to obtain their spatial distribution (>400 events from 11 tissues, Fig.4D). One intuitively would have expected that intercalations occurred within the constriction, and even more acutely near the neck of the channel, channel where the elongational component of the base flow was maximized. This was indeed the case for softest explants (distribution in Fig.4D, median x position in Fig.4E). In rigid explants, however, intercalations occurred through a broad area that covered the pre-chamber and the funnel (Fig.4D,F), consistent with stress caused by the crowding in the constriction and the channel entrance propagating at a distance. For a subset of explants, the membrane signal was sharp enough to visualise with good accuracy cell-cell junctions over the whole time of an intercalation event, starting from a broad contact until regrowth of the new junction (Fig.4A,B). We measured the length of the shirking junction until its complete disappearance, then the length of the new junction formed after the T1 transition (Fig.4C). The rate of decrease in junction length as well as the minimal length just preceding contact rupture were surprisingly homogenous, considering the huge differences in global rheology of individual explants. The rate of growth of the new junction, although more variable, was still within the range of the shrinkage rate. Intuitively one may expect that the direction of the global stress on the tissue may affect the way contacts remodelled. Surprisingly, the orientation of the T1 transition had no impact on the rate of junctional length shrinking or growth (Fig.4C’). There was a slight trend for faster remodelling in softer tissues (Fig.4C”).

**Figure 4.**
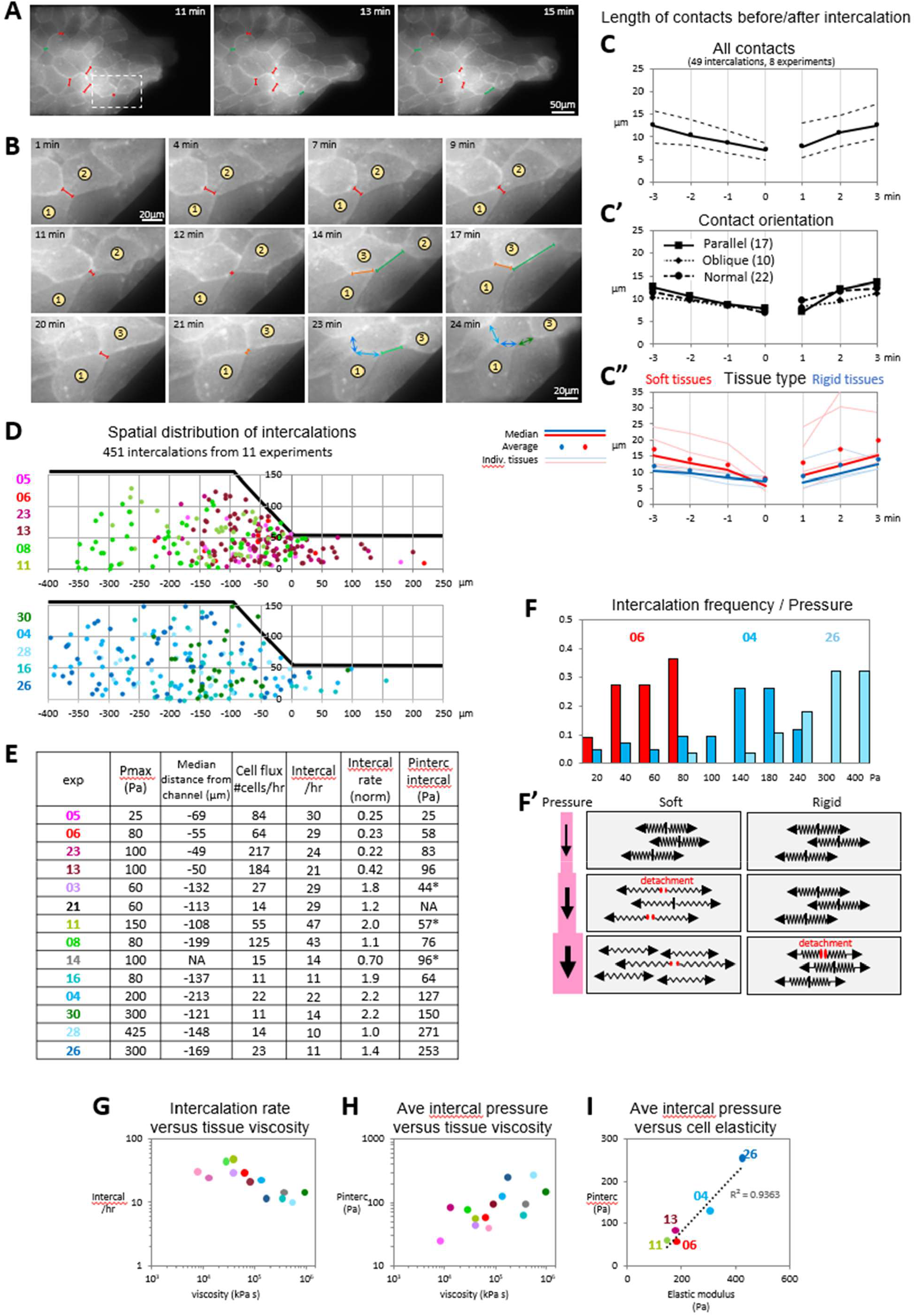
Intercalation events: Dynamics, distribution and pressure dependency. **A)** Three frames from time lapse of experiment #11 (Movie 3). The aggregate is undergoing multiple intercalations. Selected shrinking contacts are highlighted in red, new expanding contacts in green. Scale bar, 50µm. **B)** Detail from the same aggregate, showing the sequence of two intercalation events. During the first event, cell (1) separates from cell (2), in the second event cell (1) separates from cell (3). Shrinking contacts are in red or orange, new extending contact in tints of green. Between 23 and 25min, a phase of retraction occurs, which manifests in shrinking of the new contact (dark green double arrowhead) and of the adjacent contact (light to dark blue double arrowhead), both oriented in the direction of aspiration, while the contact normal to the channel expands (dark to light blue double arrowhead). Scale bar, 20µm. **C)** Kinetics of contact shrinkage and extension. Contact length was measured as shown in panels A and B. T = 0 corresponds to the last frame before T1 transition. Dashed lines indicate Q1 and Q3 quartiles. **C’)** Median values for intercalations categorized according to the orientation of the initial contact relative to the channel, i.e. normal, parallel, or oblique (45°+/-5). The three categories showed no significant difference. **E”)** Median and average values for the soft (#05, 06, 08, 11, 13, 23) and rigid tissues (#04, 16, 26, 28, 30). **D)** Spatial distribution of intercalations. The position of all discernible T1 transitions was mapped relative to the axis of the chamber (y = 0) and the entrance of the channel (x =0). On the y axis, both sides were given positive values to superimpose all events. The two maps compile data from soft and rigid tissues. In soft tissues, intercalations tended to occur close to the entrance of the channel, in rigid tissues, further back. **E)** Compilation of various quantified parameters. The table indicates, for each experiment, the maximal pressure reached, the median x distance of intercalation relative to the entrance of the channel, the cell flux, quantified as rate of cells entering the channel/hr, the number of detectable intercalations, both absolute and normalized to the global cell flux. Stiff tissues showed a significantly higher relative rate of intercalations than soft tissues. The last column indicates the average pressure Pinterc required for intercalation, estimated from the frequency of intercalations (F). Asterisks: Pinterc before tissue rupture. **F-F’)** Estimate of intercalation frequency as function of pressure. F) Distribution of intercalations at subsequent pressure steps in three representative tissues. F’) Intercalation frequency as proxy for adhesive strength: Detachment of cell-cell contacts, required for intercalation, occurs at higher pressure in rigid tissues, indicating that cell-cell adhesion is stronger. **G)** Intercalation rate (from table E) weakly inversely scales with tissue viscosity (viscosity module estimates from Figure 1). **H)** Pinterc scales with tissue viscosity. **I)** Pinterc values versus elastic modulus from Figure 2G.

A related question was the frequency of intercalations. One may expect that this frequency would be related to the viscosity of the tissue. The absolute rate of intercalations did show a slight inverse trend with viscosity, but the changes were small (Fig.4E,G). Considering the large difference in the flow rate of cells moving and entering the channel, much higher in soft tissues, we also estimated a relative intercalation rate, calculated by normalizing to the cell flow. The normalized intercalation rate turned to be lower for most soft tissues, with exceptions (Fig.4F). Thus, intercalation rates per se did not seem to directly relate to viscosity. The picture became clearer once considering the exerted pressure at which intercalations occurred: Shrinkage and disruption of the existing contact is expected to depend on the adhesion strength of the contact. In our experiments, the stress exerted by the aspiration pressure, transmitted across the elastic cells, must inevitably apply to the adhesive contacts. One can thus consider the aspiration pressure required to break a contact as readout for its strength. We reasoned that the frequency distribution of intercalations over the range of pressure exerted during an aspiration experiment could then be used to extract an estimated pondered average *“intercalation pressure*” (Pinterc) required for contact disruption, thus as a proxy to adhesion strength. The results are presented in Figure 4E and F, and the rationale underlaying this approach is graphically represented in panel F’. Pinterc correlated with the viscosity of the tissue (Fig. 4H). Importantly, Pinterc also scaled well with the elastic modulus (Fig. 4I). This latter relationship is consistent with cortical rigidity playing a crucial role in controlling adhesive contacts, as presented in Supplemental Box 2 and discussed below.

### PIV-based inference of mechanical coupling

We next used particle image velocimetry (PIV) to interrogate the global cell fluxes during aspiration. Examples of velocity maps for tissues #06 and #04 in Figure 2A illustrate the high dynamics of the tissues. One straightforward analysis consisted at correlating the average velocity of the tissue with the velocity of the front. This provided a simple readout of the underlying mechanical coupling through the tissue (Fig.5B,C,D, supplemental Fig.S3). Strikingly, moderately soft tissues #06 and #11 showed a close to perfect correlation, indicating that the stress exerted at the front was rapidly and broadly transmitted to the whole tissue. The correlation was significantly lower for the rigid tissues #04 and #28, consistent with stress dissipation due to the higher viscosity of these tissues. However, #23, one of the softest tissues in our experimental set, yielded the lowest correlation, which in this case likely resulted from the weak cohesiveness of the tissue. The same trends were observed when calculating a sliding correlation over time (grey lines in graphs of Fig.5B,C and supplemental Fig.S3). Importantly, correlation tended to increase in the late phase of aspiration (Fig.5D), indicative of reinforcement of tissue cohesiveness.

We further compared the velocity at different positions relative to the entrance of the channel. Figure 5E presents the corresponding correlations between contiguous regions. They were globally consistent with the median to front correlations. Interestingly, each profile showed a clear minimum, the position of which related to tissue properties, close to the entrance for soft tissues, increasingly far behind for rigid tissues (Fig.5D), in general agreement the distribution of intercalation events (Fig.4D,E). These velocity profiles provided additional information on tissue properties. They showed sharp accelerations that coincided with pressure increments, as well as additional asynchronous variations. The pressure-related events varied as a continuum from the most rigid to the softer tissues. Comparison of #04 and #06, respectively moderately rigid and soft tissues, highlighted key features (Fig.5B,C): The #04 profile was characterized by mild steps followed by slow relaxation, while on the contrary, the #06 profile had tall peaks soaring back to the initial baseline. Profile #11 (supplemental Fig.3) appeared intermediate, with peaks similar to #06, but also a progressive increase of the baseline. For the most rigid tissue #28, velocity remained evenly low until high pressures were reached (>60min ~ >200Pa). Similar to #04, #28 did show sharp peaks at late times/high pressure (supplemental Fig.3). Again, the softest tissue #23 behaved atypically, with variations in velocity uncorrelated to pressure steps (supplemental Fig.3). Taken together, these patterns could be interpreted as follows: The velocity steps clearly corresponded to the elastic response of the tissue to stress, with a magnitude inversely proportional to the elasticity modulus.

**Figure 5.**
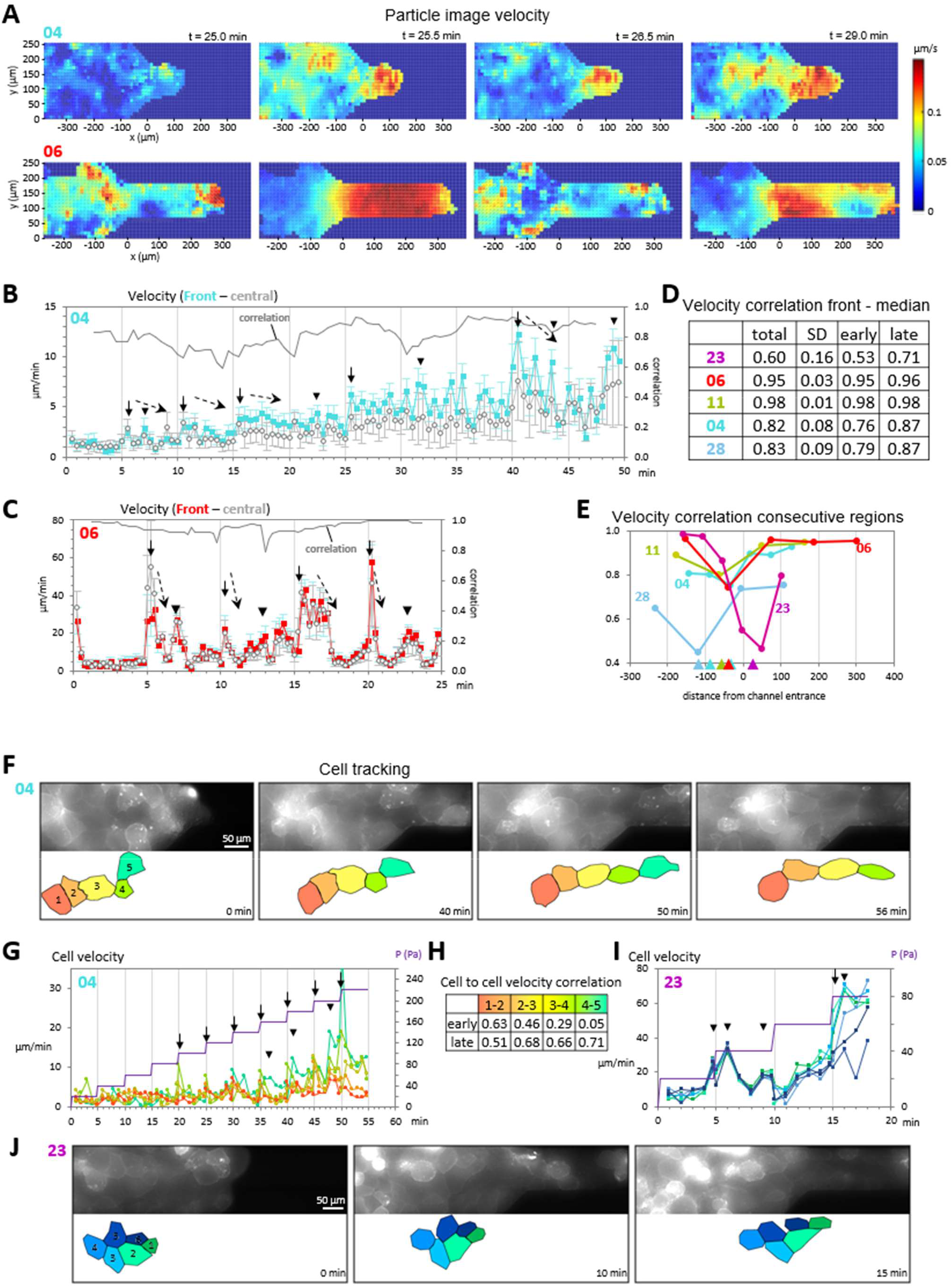
Tissue and cell velocities reflect underlying mechanical coupling. **A-E)** Particle image velocimetry (PIV). **A)** Examples of velocity fields measured by PIV for tissues #04 and 06, color-coded, illustrating high dynamics and heterogeneity. **B**,**C)** Velocities of the front edge and of the median x position of the tissue through the whole time-lapses. Vertical black arrows point to steep increases in velocity coinciding with a pressure step (vertical line, 5 min intervals). Oblique dashed arrows indicate subsequent decreases, shallow for the rigid tissue #04, fast for the soft tissue #06. Arrowheads point to velocity peaks not coinciding with pressure steps. The grey continuous line represents the sliding correlation between front and median velocities. The correlation is higher in soft tissue #06 than tissue #04. Profiles for three additional tissues are presented in Supplemental Figure S6. Error bars, SD. **D)** Summary of velocity correlations for the whole time-lapses, as well as for the first and second halves. **E)** Velocity correlation versus x position. Average velocity values were calculated for five subregions, from the back of the chamber to the channel entry and to inside the channel (see Supplemental Figure S6) and correlation was calculated between each consecutive subregion. Note a clear trend for the position of lowest correlation relative to tissue properties (arrowheads, color-coded for each tissue), farthest from the channel for the most rigid tissues, near the channel entrance for soft tissues. **F-J)** Velocity of single cells tracked along the time-lapse of tissues #04 and #23. **F**,**J)** Selected frames and corresponding schemes of segmented cells. Velocity peaking with pressure steps (vertical arrows) are prominent in #04, while #23 showed mostly unrelated peaks, consistent with PIV data. **G, I)** Corresponding velocity profiles. **H)** Cell to cell velocity correlations during the first and second half of the movie for tissue #04. In all but the rear cell pair, correlation increased at late time points, consistent with cell-cell coupling as cells aligned and stretched (F).

The subsequent decay reflected the ability of the tissue to adapt to increased stress through cell rearrangement. Fast extensive decay in tissues #6 and #11 was consistent with intercalation events readily occurring at low pressure (low adhesive strength), as opposed to stiff tissues #04 and #28, where high adhesion restrained intercalation until high pressures were reached. As for peaks that appeared unrelated to pressure increments, they likely reflected the autonomous and heterogenous activities occurring within the tissue.

To relate PIV data to single cell behaviour, we tracked small groups of cells, and calculated the velocity of the individual cells. Figure 5F shows frames of a row of 5 cells from tissue #04, and Fig. 5G the plot of their respective velocities. The cell velocity profiles were close to identical to the PIV profiles (Fig.5B and Fig.S3), with increasingly strong peaks as pressure increased, interspersed with additional peaks. Consistently, pairwise correlation (Fig.5H) indicated that cell-cell coupling increased at higher pressures, consistent with the trend of PIV correlation (Fig. 5D). Fig.5I shows the profile for a group of 6 cells from the soft tissue #23, in which peaks did not coincide with pressure steps, again consistent with the PIV data. However, these two cases were particularly simple, notably lacking intercalation events. We will see below that more complex behaviours could be characterized.

In summary, this approach of velocity correlation revealed that tissues reacted to increased stress through mechanical coupling, consistent with the capacity of reinforcement of cell adhesive contacts.

### Stepping down to the cell scale: Inferences from cell contact geometry and dynamics

The shape and dynamics of cell interfaces reflect the underlying mechanics (see Supplemental Boxes 1 and 2). A global glance at the mesoderm explants showed that cells of soft tissues tended to have smoother, curved contact interfaces, those of rigid tissues a more angular morphology (Supplemental Fig.S4). However, all tissues appeared highly coherent as judged by their extensive contacts, both under low pressure, or when stretched under higher pressure (Supplemental Fig.4). Thus, differences in cell morphology coincided with their relative rigidity but not necessarily with tissue cohesiveness (as defined in Supplemental Box 1). This important observation indicates that mesoderm can maintain the right balance of cortical tension and adhesion to remain coherent through a large spectrum of physical properties.

Beyond these general trends, a closer look at individual cells within local groups showed striking features. Despite the complexity of the system, some characteristic cell behaviours emerged that could be interpreted by simple considerations on cell and contact morphology (Supplemental Box 2). Supplemental Figure S5, illustrates this approach for a group of cells in tissue #11 located near the channel entrance. Most events could be dissected into basic processes (Supplemental Fig.5A’). Two rows of cells, color-coded respectively with shades of blue and shades of orange-red, illustrate well the dynamics of mechanical coupling, evident from their collective movement and synchronous stretching, and validated by velocity correlation (Supplemental Fig.5B,C, Movie 4). A detailed description of these dynamics is presented in the legend of Supplemental Fig.S5. Briefly, salient features of the blue row were its initial coupling, temporarily disrupted by a cell-cell detachment, then rescued through contribution of an additional cell inserted through vertical intercalation. As for the orange row, coupling at the front was interrupted by the lateral intercalation of the red cell, which exhibited a typical sliding movement (see below).

Looking now at cell contact geometry provided a readout that could at least partly predict among possible outcomes, whether detachment, with or without previous stretching, or on the contrary reinforcement, consistent with the model presented in Supplemental Box 2. Examples are detailed in supplemental Figure S6 and Movie 5. Strikingly, alignment of the lateral edges of two adjacent cells would often coincide with coordinated stretching and contact resistance, consistent with mechanical coupling. On the contrary, contacts showing a deeper indent, thus a more acute vertex angle, would rather seem more prone to rupture. These contact configurations accounted for the phases of coupling/detachment in the blue and orange rows in Supplemental Fig.S5. High sensitivity of cell contacts to shearing (Kale et al., 2018) appears to account for frequent situations of cells sliding relative to each other. Such phases were often concluded by transition to mechanical coupling that systematically coincided with cell edges becoming aligned (Supplemental Fig.S5, red cell 1080s and supplemental Fig.S6A,B). Such behaviour is suggestive of a phenomenon of adhesion “clamping”, which will deserve further investigation in the future. A related remarkable phenomenon specifically took place along the wall of the constriction, which we named the “sliding slug phenotype”, where cells became roundish and slithered along the corner of the constriction, eventually recovering a normal shape and contacts once inside the channel (supplemental Fig.S5 and S6C,D, Movie 4). As highest shearing happens to be predicted to occur close to the channel walls, this slug phenotype, although resulting from the artificial configuration of the microchip, nicely highlights the effect of shearing forces on cell adhesion.

### Active protrusive behaviour

So far, we have addressed the tensile and adhesive aspects of the system. Another important aspect is the ability of cells to emit protrusions, which interact not only with the matrix, but also with other cells. In our settings, only the latter were relevant, since the physical substrate was non-adhesive. In bright field movies, intense activity at cells borders could be detected. We thus performed high speed imaging using spinning confocal microscopy. We took advantage of the mosaic expression of membrane-targeted fluorescent protein (here mCherry) in these reaggregated tissues and focused on bright cells well resolved from a background of paler/unlabelled cells (Supplemental Fig.S7). Most cells showed indeed extensive protrusions spreading on neighbours, best viewed at the bottom plane of the tissue (Supplemental Fig.S7A, Movie 6). Importantly, these protrusions were unequivocally actively extending, and could be unambiguously distinguished from retraction fibers that were passively stretched as cells pulled on each other (Supplemental Fig.S7A). Protrusions were found both at the front of the cells (in the direction of the aspiration) and at the rear, thus cells were actively reaching for both leader and follower cells. More than 75% of the analysed cells showed active protrusive activity, with almost half displaying both forward and rearward protrusions (Supplemental Fig.S7B). We looked at potential correlation between cell displacement speed along the channel axis and the presence of protrusions. Cells forming rear protrusions (with or without front protrusions, supplemental Fig.S7C) had a significantly lower speed than other cells from the same tissues, consistent with this process contributing to active resistance against the flow. There was no clear correlation between forward protrusions and cell dynamics.

## Discussion

We used here a minimalistic synthetic system to explore the diversity of cell behaviour taking place in the highly dynamic mesoderm tissue, and to relate cell-scale events to global tissue rheology. Keeping in mind the semi-quantitative nature of our analysis, it does provide a coherent picture of remodelling of a compact mesenchymal tissue and highlights general properties that emerge from beneath the tissue complexity and the embryo to embryo mechanical variability. Firstly, the mesoderm manifests a striking cohesiveness irrespective of this large variability. Importantly, it appears to be able to remain cohesive even under high stress as imposed here by pressure increments, cells maintaining extensive cell-cell contacts even when severely stretched. As a matter of fact, many identified features of collective behaviour appeared to be shared by soft and rigid tissue, the major difference being that they occurred at different pressure regimes. It thus appears that mesoderm cells are capable of remarkable adaptability when mechanically challenged, both in terms of cell stiffness and cell adhesive strength. The reinforcing effect of cell tension on adhesion of mesoderm cells is flagrant in our movies: Stress applied by aspiration translates in a characteristic reaction that includes, in addition to cell stretching, unambiguous cell-cell coupling, reflected by high displacement correlation of adjacent cells and resistance of the contact to detachment. The phenomenon is observed in both soft and stiff samples, albeit to different degrees, and obviously at different levels of pressure. These observations are consistent with the fact that beyond its antagonistic effect on cell adhesiveness (Supplemental Box 1), increased cortical tension can stimulate “link adhesion”, ultimately reinforcing adhesive bonds (Maître and Heisenberg, 2013; Winklbauer, 2015)(Supplemental Boxes 1 and 2). This facet appears to be surprisingly salient in this mesoderm tissue.

Figure 8 presents a simple model that attempts to recapitulate these major observations. Our data indicate that cell stiffness, aka cortical contractility, is likely to be the major parameter that accounts for the tissue properties. Differences in cell deformation in response to stress not only readily account for the corresponding differences in tissue elasticity, but the cell elastic modulus also scales well with adhesive strength (Pinterc), thus ultimately tissue viscosity. This is well in line with the notion that the apparent viscosity of a tissue reflects resistance to cell-cell rearrangements. The stronger contact resistance in rigid tissues is also consistent with a broader distribution of intercalations compared to soft tissues, where cell rearrangement readily occurs near the channel entry, where constrains are highest (Fig.8B). Additionally, the slower pace of aspiration in stiff tissues may have given more time for intrinsic cell activity leading to spontaneous intercalations, thus less directly dependent of constrains imposed by the constriction.

Cell morphology analysis, even though performed on a limited number of cells, revealed geometry of cell interfaces as a surprisingly good predictor of cell behaviour (Fig.8C,D), supporting a model where contact detachment or reinforcement depends on both strength and orientation of applied tension. This model is consistent with adhesive contacts being much more resistant to normal forces than to lateral or shearing forces (Evans, 1985; Garrivier et al., 2002; Kale et al., 2018; Rozema et al., 2025)(Supplemental Box 2). Our previous work had distinguished two antagonistic regimes for cadherin contacts, peeling versus condensation, which depended precisely on the orientation of tension exerted by the cell cortex. The present study complements this picture by highlighting another frequent configuration where cell edges did align with the orientation of stress, which coincided with increased mechanical coupling (Fig.6C), reminiscent of the well-known properties of adherens junctions of epithelial cells (Arora et al., 2025; Charras and Yap, 2018; Eckert et al., 2025). The second remarkable behaviour was lateral sliding, which on the contrary related to decreased coupling, consistent with adhesion weakening under shear (Fig.6D). The latter behaviour was most acute in the form of the slug-like behaviour occurring at the entrance of the channel, the predicted site of highest shear. Strikingly, coupling was almost instantaneous recovered once propitious conditions appeared to be met, typically when the edges of two cells became aligned, which appeared to “clamp” the contact, resulting in transition from sliding to coupling (Fig. 8D).

**Figure 6.**
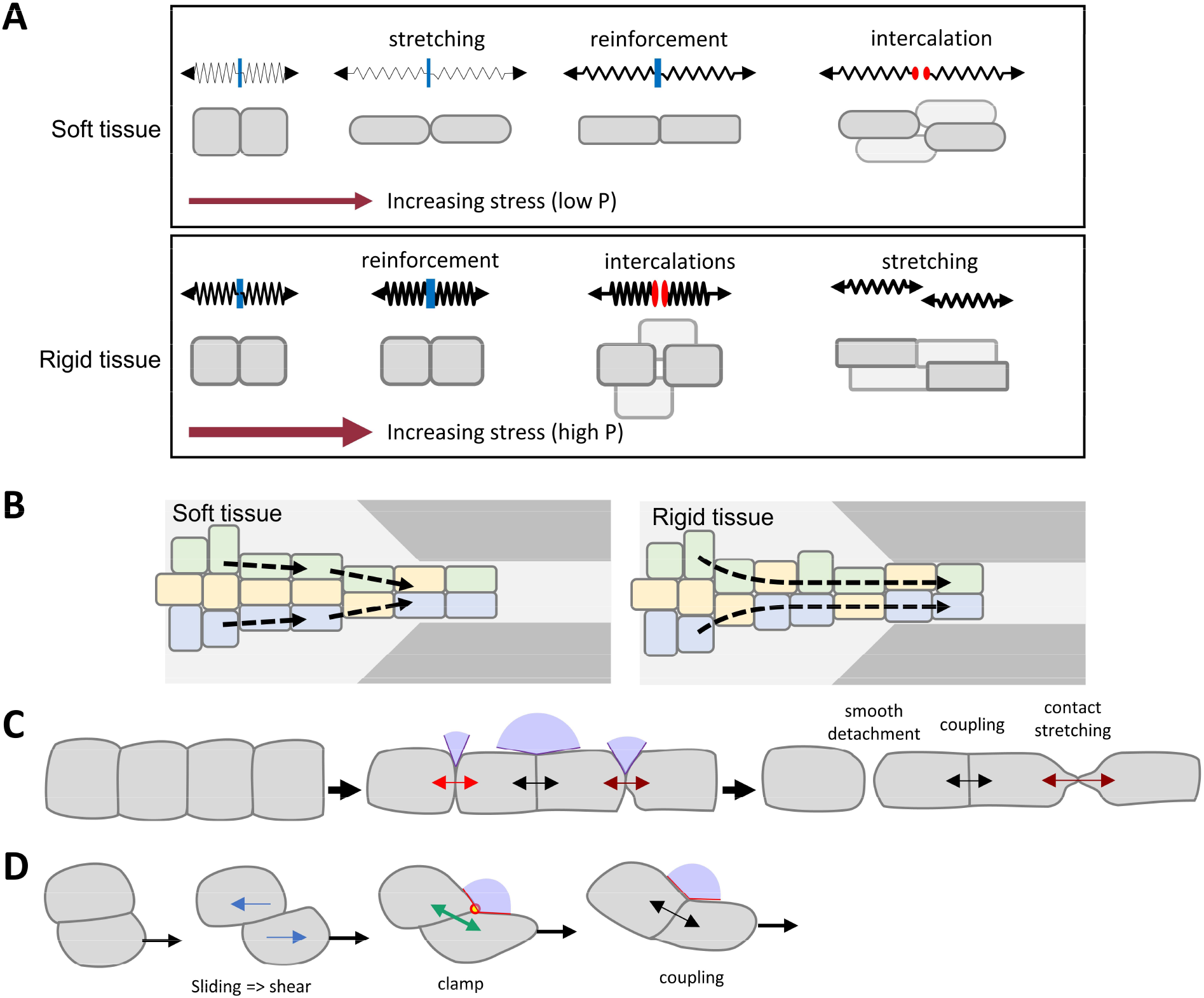
Summary model. **A)** Upon stress, the elastic response causes cells to stretch to various degrees, while at the same time, tension challenges cell-cell contacts. Under specific configurations (see C,D), cells actively react through reinforcement of cortical tension and adhesion, resulting in increased cell-cell coupling. As tension increases, it eventually overcomes the resistance of the contact, favouring detachment, and cell-cell intercalation. Soft tissues may be considerably stretched before hitting this point. In a rigid tissue, higher forces are required to deform cells, and although the system can also reinforce, the point of contact rupture may be reached earlier relative to cell elongation. **(B)** In the context of the channel constriction of aspiration experiments, soft tissues can extensively deform thanks to low cell elasticity. Intercalations then concentrate near the entrance of the channel, as response to highest geometry constraint. In rigid tissue, the higher resistance of cell cortex limits deformation, thus stress can act directly on cell contacts at a distance from the channel entrance, thus accounting for the more widespread distribution of intercalations. **(C)** Response of cell contacts to stress varies, depending in particular on the balance between contact tension and cell cortex tension. The geometry of the contact is predictive of the behaviour (Supplemental Box 2). A narrow contact angle is generally permissive for a smooth detachment. An intermediate angle rather favours contact stretching before rupture. Contacts displaying a wide angle provide a configuration where adhesion at cell-cell vertices aligns with the lateral cortex, optimal for high coupling through both cell stiffening and adhesion reinforcement. These various behaviours can be observed in soft and rigid tissues, depending on the pressure applied. **(D)** Another characteristic phenomenon is that of cells sliding over each other, consistent with adhesion poorly resisting shearing forces. However, this low adhesion regime is generally short lived. As a contact angles happen to widen, a transition evocative of clamping results into cell-cell alignment and mechanical coupling.

It will be important in future work to investigate in detail the cellular and molecular organization underlying adhesive coupling between mesoderm/mesenchymal cells and uncover the mechanism of the presumed clamp mechanism. Protrusive activity is another aspect that needs to be integrated in the picture. Our preliminary observations of protrusions forming seemingly independently of applied stress illustrate the strong intrinsic activity of these cells, but to what extent this protrusive activity contributes to tissue rheology is a non-trivial question to address.

The large physical variability between embryo batches was one of the aspects that we were keen to explore. Although a variety of cases were observed, with outliers present for any parameter, some general trends could be drawn. Tissue viscosity and elastic modulus roughly scaled together, which suggested that in some way both aspects of the tissues are related. One first piece of answer is the scaling of adhesive strength with the apparent cell elastic modulus (Fig. 4I), a strikingly property indicative of a direct link between cortical tension and adhesive strength (Winklbauer, 2015). As discussed above, this is consistent with cortical tension playing a major role of adhesive reinforcement in these tissues. On the contrary, cadherin levels do not seem to have a major impact on gastrulation (Ninomiya et al., 2012). Thus, the well-known highly variable cortex rigidity/contractility may turn out to be a simple explanation for the natural variability in tissue rheology in these early embryos.

### Study limitations

The standard tissue rheology model used here only roughly approximates the complex active properties of the tissue. Furthermore, tissue adaptation to applied forces strongly affect its rheology (David et al., 2009). One should thus consider the calculated values are mere trends. Our analysis at the cell scale also remains highly simplified. For instance, the tissue being 2-3 layer thick, cell behaviour in the lower layer examined here were subjected to unknown interactions with cells positioned above. In principle, it would also be important to perform a parameter exploration by experimentally tuning contractility and adhesion. However, we are currently missing tools that would specifically target one function, as so far it has not been possible to dissociate these intricately interconnected cell activities.

## Materials and Methods

### Embryo injection, tissue isolation and cell dissociation

Xenopus laevis used in this study were acquired from Nasco, USA, or TEFOR Paris-Saclay, France. All Xenopus research performed for this study was approved by the “Direction Generale de la Recherche et l’Innovation”. Embryos were injected equatorially in the two dorsal blastomeres of the four-cell stage embryo with in vitro transcribed mRNA coding for membrane-targeted GFP or Cherry (200 pg mRNA) as described (Kashkooli et al., 2021). Prechordal mesoderm was dissected at stage 11. Dissections were performed in 1X MBSH (88mM NaCl, 1mM KCl, 2.4 mM NaHCO_3_, 0.82 mM MgSO_4_, 0.33 mM Ca(NO3)_2_, 10 Mm Hepes, and 10 ug/uL Streptomycin and Penicillin, pH 7.4). Mesoderm explants were dissociated into single by incubation for 5-10 min in alkaline dissociation buffer (88 mM NaCl, 1 mM KCl, and 10 mM NaHCO_3_, pH 9.5).

### Microfluidic chip design, aggregate formation and aspiration protocol

The experiments were performed on a microfluidic chip made of PDMS. The PDMS mould was made by using standard lithography techniques. The microfluidic chip design is presented in Fig.1A. The whole system had an even height of 100 µm. The pre-constriction and post-constriction chambers had a width of 250 µm, the funnel-like constrictions were at a 45° angle, leading to the 100µm-wide channel. Before the experiment, all channels were passivated with 5 mg/ml of bovine serum albumin to prevent unspecific adhesion to the channel sides or bottom glass. A loading funnel made of a cut 200 µl pipette cone plunged in the microfluidic reservoir. The other side of the microfluidic chip was connected to a reservoir tube. The entire system was filled with 1X MBSH. The height difference between the microfluidic chip and the reservoir tube determined the “zero pressure”, which was obtained by adjusting the pressure to obtain a null flow inside the chip. All experiments were at 23°C. Dissociated were loaded and let sediment in the reservoir, where they started to reaggregate. A compact tissue was formed within 1hr. It was slowly aspirated in the chamber and the constriction until the entrance of the channel. Pressure was set back to zero, adjusted if required, and monitored over several minutes to verify its stability. The aspiration experiment started with 5min at P=0Pa, followed by stepwise pressure increments of 20Pa every 5 minutes. For some experiments, tissues appeared to approach a breaking point, in which case the aspiration pressure profile was adapted by extending steps to prevent the tissue rupture. Tissues were imaged every 15sec with an epifluorescence inverted Zeiss Axiovert microscope with a 20X objective (NA 0.75), equipped with an OrcaFlash4.0LT Hamamatsu CCD camera. Spinning dick confocal imaging was performed on an Andor Dragonfly equipped with dual iXon888 EMCCD cameras and controlled by Fusion (Andor), using a 20X 0.75 NA objective (Nikon).

### Image analysis

Whole tissue deformation along the channel axis was obtained by measuring the advance of its front, extracted through simple thresholding using ImageJ. Fitting curves and elastic moduli / viscosity calculations using modified Kelvin-Voigt viscoelastic model were realized on Matlab. The velocity field of the tissues was computed using 2D time-resolved PIV. Note however that no particles were added to the tissue and that the contrast provided by the cell contours was sufficient to apply the image cross-correlation algorithm. The open source PIVLab software was used to perform adaptative cross-correlation of the images with an overlap between adjoining interrogation areas set to 50%. The average tissue velocity corresponded to the average within the central longitudinal region of the channel (in order to exclude the regions next to the channel walls). All manually cell segmentation, as well as measurements of contact length and orientation, and ellipse fitting were performed using ImageJ.

## Supporting information

Supplemental Data (Table, Boxes 1+2 and Figures S1-S6)

## Acknowledgments

We acknowledge the imaging platform MRI (Montpellier Ressources Imagerie) for providing access to microscopy facilities and technical support. We also thank the animal facility ZEFIX (zebrafish and Xenopus platform) for maintaining our animal colonies and providing expert care. This work was supported by the ANR (Agence Nationale de la Recherche) grants ICM (ANR-14-ACHN-0004) and Inters-cal (ANR-21-CE13-0042) to F.F. and Lovetiss (ANR-21-CE06-0023–01) to L.C., and from the Labex NUMEV (ANR-10-LABX-0020) within the I-Site MUSE to L.C.

