## Supplemental Data (Table, Boxes 1+2 and Figures S1-S6) for "A microfluidic approach to explore mesoderm tissue dynamics and its natural variability"

### Box 1

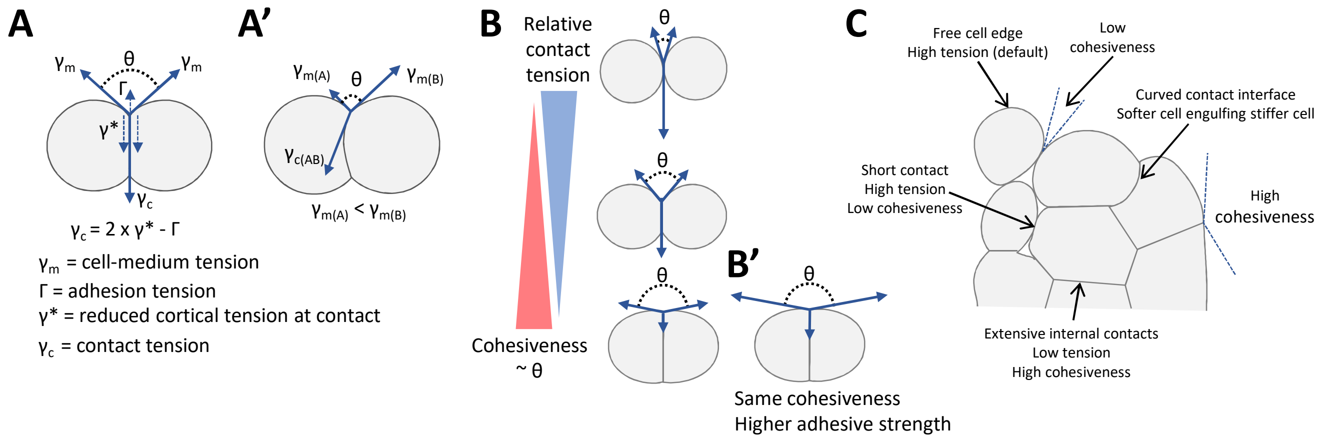


**Box 1: Cell contact geometry, interfacial tension and adhesiveness**

Cell and tissue geometry reflect the underlying mechanics. Cell-cell adhesion at a contact is modelled as the balance between cortical contractility and cell adhesion (Brodland 2002, Lecuit and Lenne, 2007, Winklbauer 2015). **(A)**. This balance is typically represented by the equilibrium of forces acting at a contact vertex. In the simplest configuration of a cell doublet, the vertex is the corner point where the cell-cell contact meets with the two "free edges". Free edges (or cell-medium interfaces) are under tension $\gamma_m$, mainly built by the actomyosin cell cortex, thus typically called *cortical tension*. At the contact interface, several tensions are at play, with the contribution of the cortical tension γ* of both cells, as well as the adhesive energy produced by engagement of adhesion molecules, which corresponds to a negative tension Γ. Cell *contact tension* $\gamma_c$ is defined as the sum of these three tensions ($2 \times \gamma^* - \Gamma$). Importantly, the major effect of cadherin adhesion is generally not the adhesive energy alone, but rather a local downregulation of cortical tension along the adhesive contact ($\gamma^* < \gamma_m$). **(A')** The same balance of tensions also applies to the case of cells with different cortical tension ($\gamma_{m(A)} < \gamma_{m(B)}$) (Canty et al, 2017). Here the cell contact is curved, the softest cell tends to engulf the stiffest. **(B)** The degree of *cohesiveness (adhesiveness* in Parent et al, 2017) is a value ranging from 0 to 1, which reflects the ratio between contact tension and free cortical tension, and directly relates to the angle θ formed by the two edges at the vertex. **(B')** Importantly, the same *cohesiveness* can be achieved for softer and stiffer cells, provided that $\gamma_c$ is proportionally reduced. However, for the same *cohesiveness*, adhesive contacts of stiffer cells can withstand higher *absolute tension* (Winklbauer 2015), thus display a higher effective *adhesive strength*. **(C)** The model applies to tissues, allowing *force inference* based on cell geometry. When cells are highly adhesive, they maximally extend contacts, resulting in a highly *cohesive tissue.* The smaller contacts in a loose tissue are consistent with low adhesiveness. As in doublets, curved interfaces in a tissue reflect tension differences between cells.

### Box 2

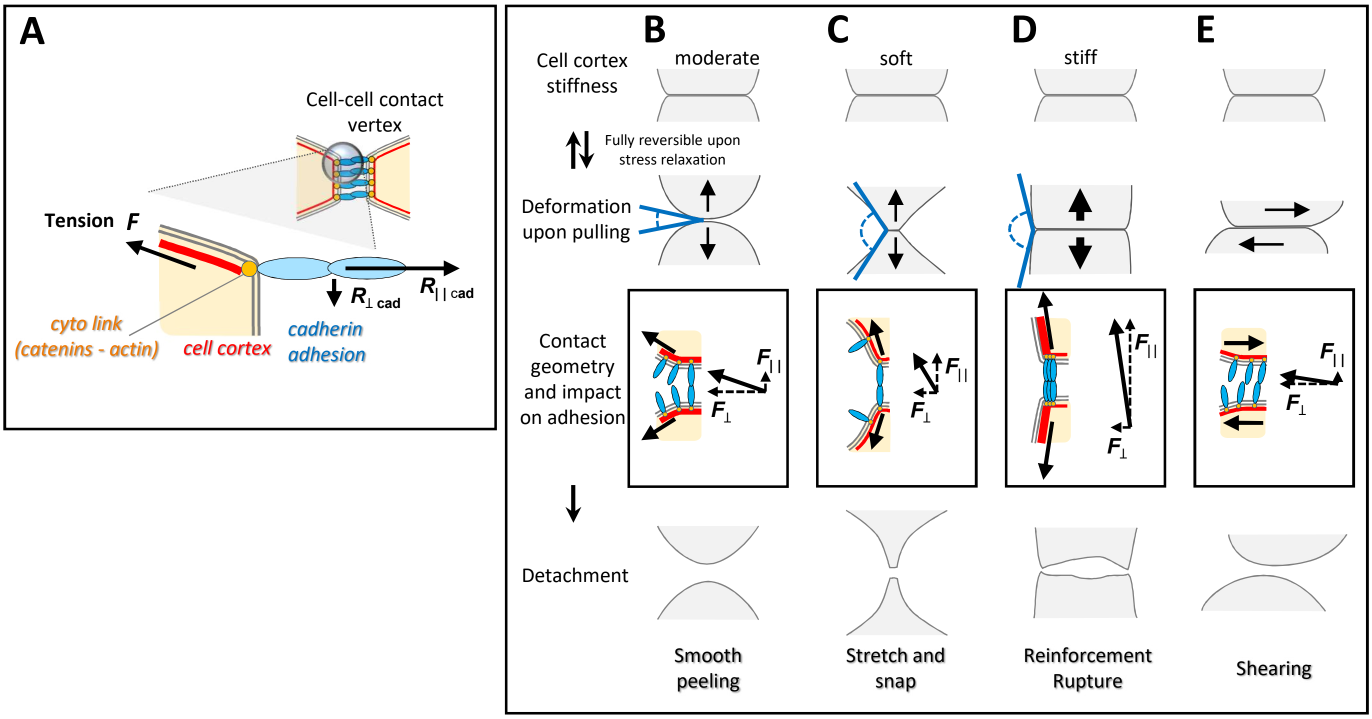


**Box 2. Contact remodelling in mesenchymal cells: Impact of tension and geometry**

The model based on Rozema et al (2025) poses that contact disassembly is dictated by a few simple rules: (**A**) Tension ***F*** exerted on a cell propagates mainly along the lateral cortex toward the cell contact vertex. The response of cadherin adhesions to stress is highly sensitive to geometry. Cadherin adhesive interactions are most resistant to forces aligned with the cadherin-cadherin axis ($\boldsymbol{R}_{|\,|\ \mathrm{cad}}$) but easily destabilized by orthogonal forces ($\boldsymbol{R}_{\perp\ \mathrm{cad}}$). (**B-E**) Typical modes of contact disassembly. A contact will differently react to stress exerted normal to the contact depending on the orientation of the tension, which in turn depends at least partly on the stiffness of the cortex. (**B**) For a moderately stiff cortex, the contact will tend to progressively open maintaining a narrow angle, mostly impacting on $\boldsymbol{R}_{\perp\ \mathrm{cad}}$. This favours a smooth detachment through cadherin "peeling" and lateral diffusion. (**C**) A soft cortex will easily deform, increasing the $\boldsymbol{R}_{\perp\ \mathrm{cad}}$ component. This results in transient resistance, but the weak cortex stretches and the contact eventually snaps. (**D**) A stiffer cortex can resist deformation. With stress at the vertex oriented along the axis of cadherin highest resistance, cadherin-cortex coupling can be stimulated, reinforcing adhesion. Here stronger tension will be required to rupture of the contact. (**E**) A different situation is achieved when two cells, rather than pulling away normal to the contact, slide relative to each other. Here shearing forces are predicted to efficiently destabilize cadherin adhesion, as show by Kale and colleagues (2018). For cells in the context of a complex tissue, each contact will remodel according to the combined influence of basal cell/cortical stiffness, geometry, as well as intensity of the exerted tension.

**Table 1.** Comparison of tissue scale and cell scale values between this study and previous reports. David et al, 2014; (2) Canty et al, 2017; (3) single cells in suspension; (4) Cited from Marmottant 2009.

| | This study | Previous (Xenopus) | Tlili et al 2022 (carcinoma F9) |
|---|---|---|---|
| Tissue scale viscosity (Pa.s) | $10^4$-$10^6$ | 5-50x$10^3$ (1) | $10^5$ (4) |
| Tissue scale elastic modulus (Pa) | $10^2$-$10^4$ | ND | ND |
| Tissue scale relaxation time (s) | 10-$10^3$ | ND | $10^3$ |
| Cell scale elastic modulus (Pa) | 100-400 | 200-1000 (2)(3) | $10^2$ |

### Figure S1

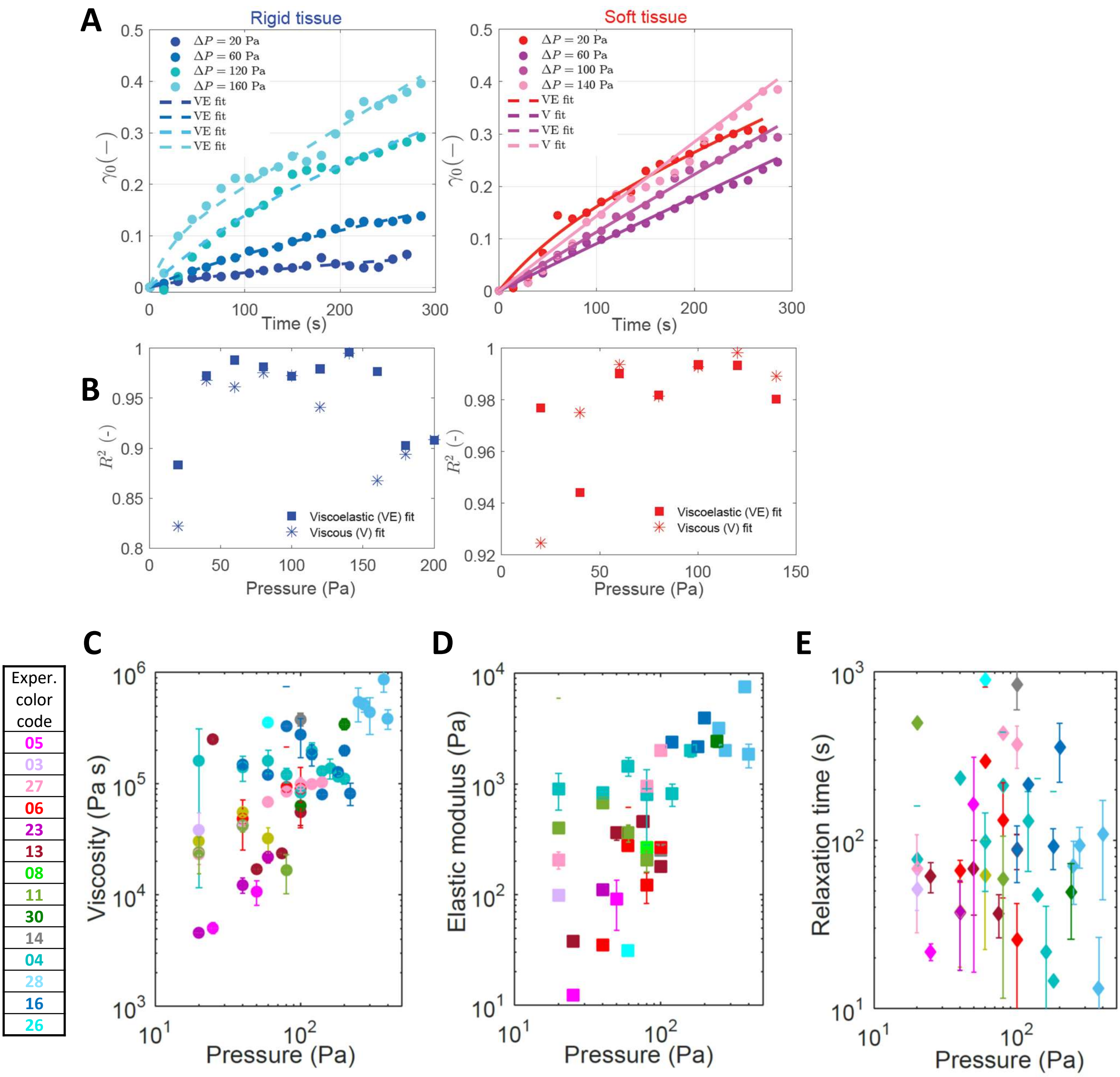


**Aspiration curve fitting and rheology values.** Each experiment is color-coded. For each pressure step, the advancement (l) of the explant inside the constriction was tracked and computed its deformation (strain) computed as $\gamma = (l - l_0)/w$, where $l_0$ is the initial position for each step and $w$ is the width of the thin channel. Each step was fitted using a modified Kelvin-Voigt viscoelastic model which predicts a temporal evolution of the strain given by $\gamma = \gamma_0\left(1 - e^{-(t-t_0)/\lambda}\right) + \dot{\gamma}_\infty(t - t_0)$, where $\gamma_0$ is the intercept, $\dot{\gamma}_\infty$ is the slope of the linear phase and $t_0$ corresponds to the initial time of each step. The first term of the equation accounts for the fast elastic increase observed at short times and the second to the viscous behaviour observed at longer times. The relaxation time ($\lambda$) qualitatively sets the transition between the two different flow behaviours. This equation has three fitting parameters ($\dot{\gamma}_\infty, \gamma_0, \lambda$) and it is difficult to have a unique set of resulting parameters. In order to improve the fitting procedure, we first fitted the last part of each step with a linear fit ($\gamma = \dot{\gamma}_\infty(t - t_0)$) corresponding to the viscous phase, which was easier to fit robustly, and we used the resulting value obtained for $\dot{\gamma}_\infty$ as an initial starting parameter to optimize the viscoelastic fit over the entire step duration. For some experiments, only one or two segments were fitted, others had up to 10 segments. **A)** Illustration of fitting for two examples, of a rigid and a soft tissue. In each case 4 curve segments are shown, with the corresponding applied pressure. VE and V stand for viscoelastic and viscous fit, respectively. For each step both fits were performed and the fit providing a R2 closer to 1 was selected. **B)** R2 values for fitting of each curve segment. Only fits with R2 above 0.8 were considered. Fits with lower R2 values were excluded for the rheological analysis. Asterisks indicate values from fitting with a purely viscous model. **C,D,E)** Estimates of viscosity modulus, elastic modulus and relaxation time. The error bars correspond to the confidence intervals provided by the fitting procedure.

### Figure S2

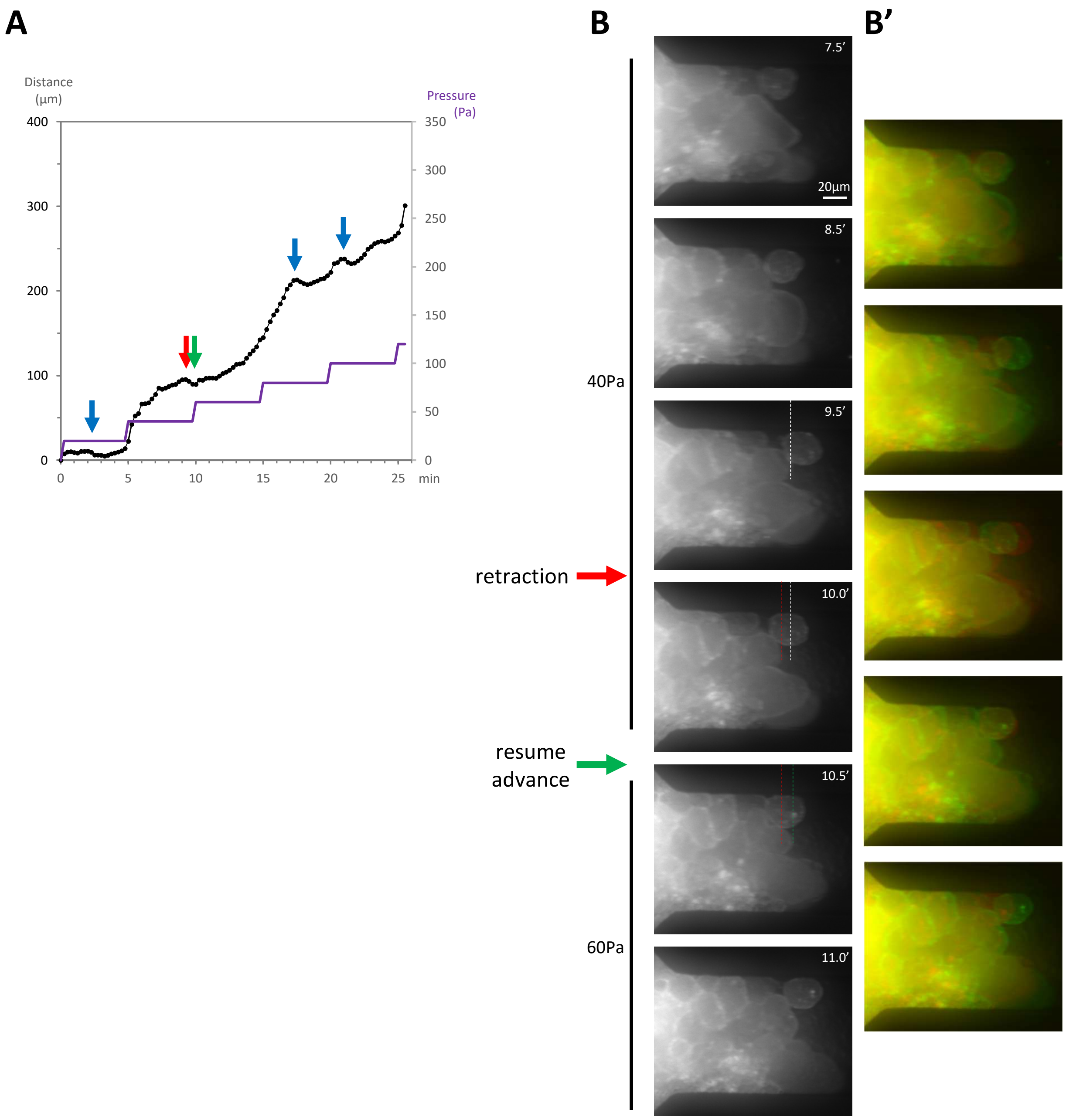


**Supplemental Figure S2. Example of abrupt tissue retraction.** From time lapse movie of experiment 06. **A)** Aspiration profile. Arrows point to retraction events. **B)** Cropped region of the entry of the channel. The aggregate steadily advanced under constant 40Pa aspiration, until 9.5min, at which point it abruptly retracted (red arrow). Advance was restored less 1min later. In this case it coincided with a pressure step to 60Pa, but retractions followed by re-elongation also occurred independently of pressure (blue arrows). Dashed white, red and green lines serve as benchmarks, **B’)** Superimposition of two successive frames, the first showed in red, the second in green.

### Figure S3

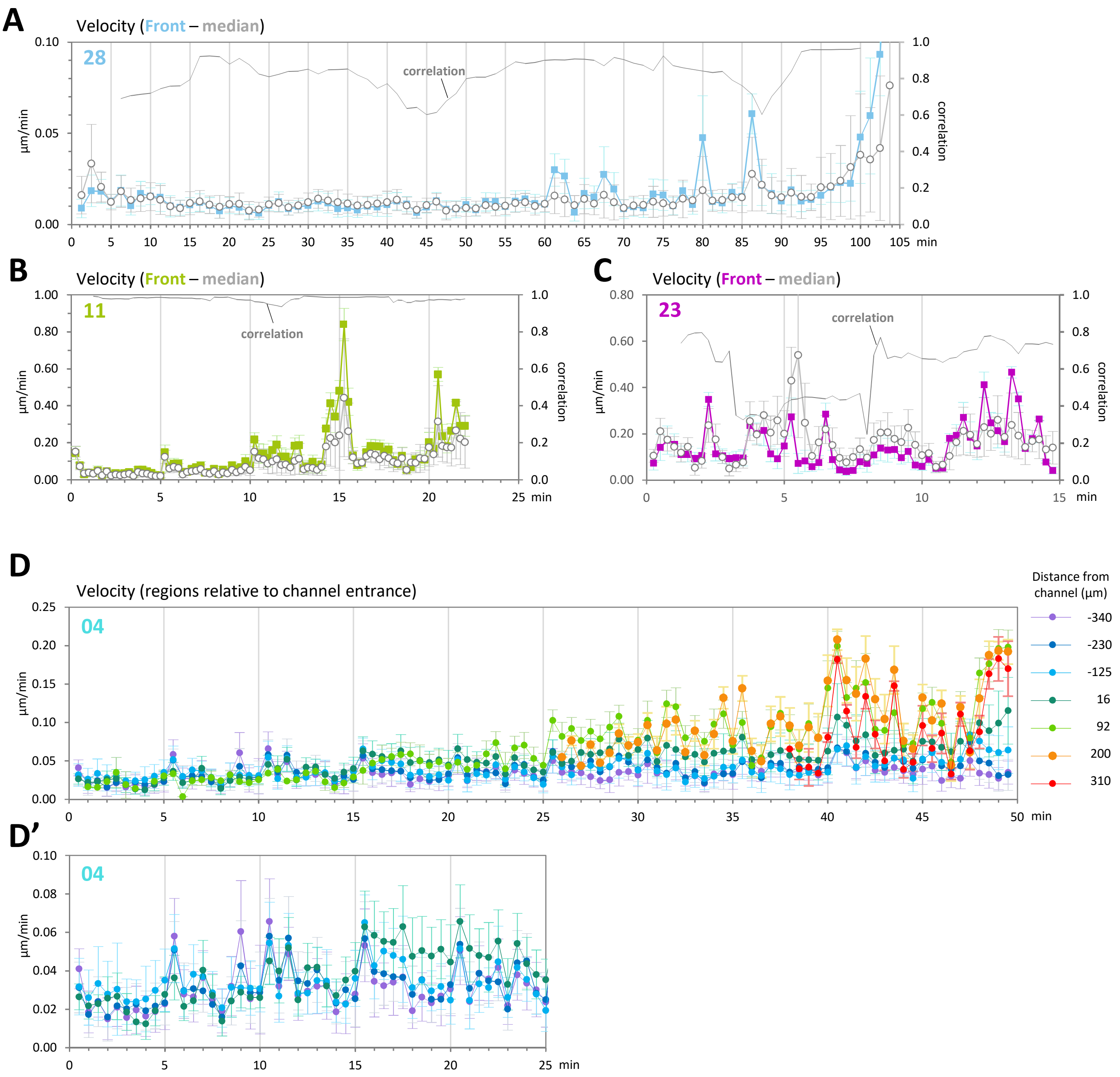


**Supplemental Figure S3.** Tissue velocity for tissues #28, #11 and #23, corresponding respectively to the most rigid, intermediate and softest tissues. **A-C)** Velocities for front and for median position of the tissue, and corresponding correlation. **D)** Velocity in consecutive regions along the x axis, relative to the entrance of the channel. **D')** First half of the graph, adjusted scale. Error bars, SD.

### Figure S4

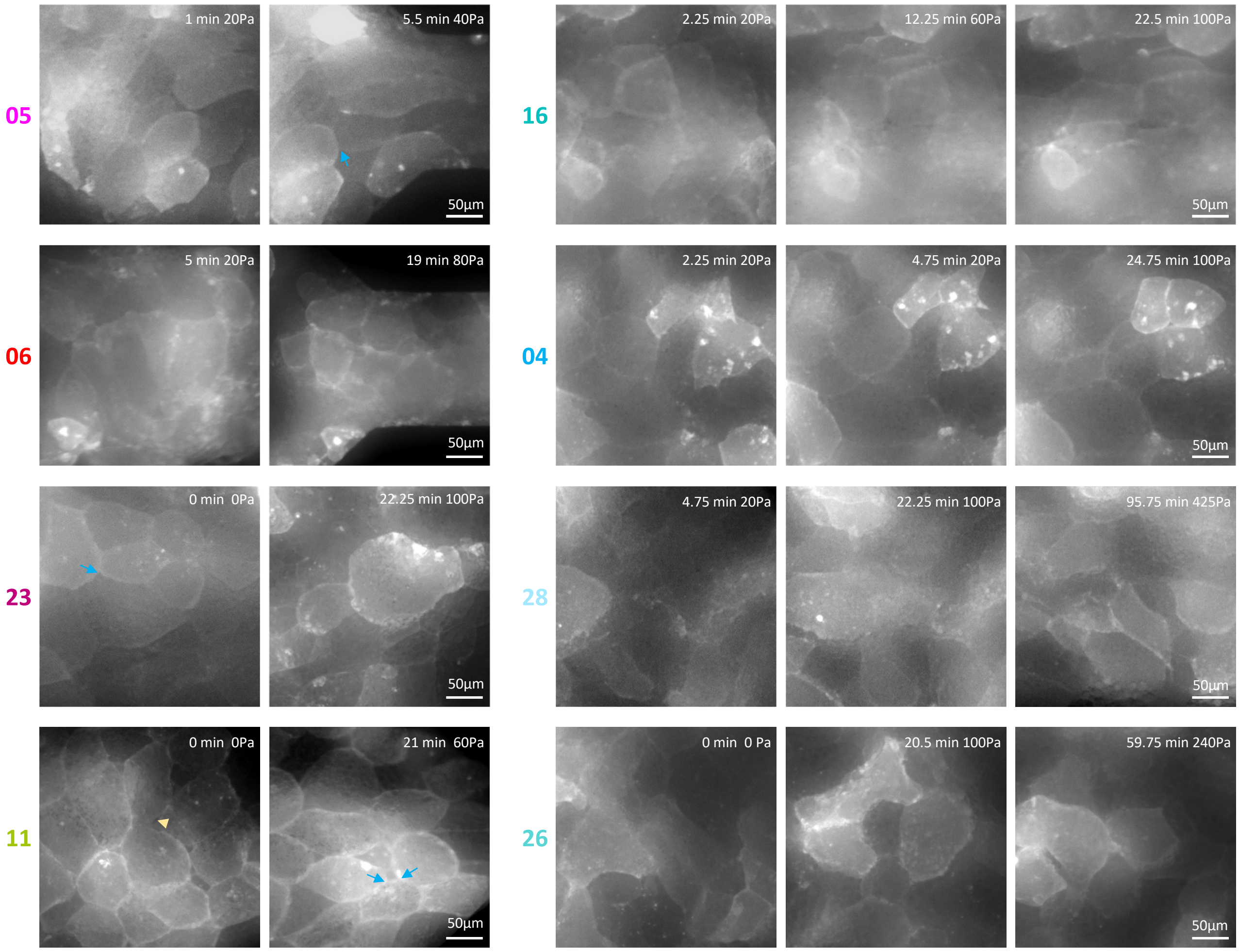


**Supplemental Figure S4. General cell morphology in tissue explants.** Details from frames of movies for soft (left) and rigid (right) tissues, at early time point under no/low pressure (0-5min), and a later time points during aspiration. While heterogeneous, cell morphology is clearly more roundish in soft explants, and more polygonal with more marked vertices in rigid tissues. However, even in soft tissues where cell edges are curved and/or cells strongly deformed, cell contacts are extensive and cell vertices are rather tight (blue arrows) except for rare cases (yellow arrowhead).

### Figure S5

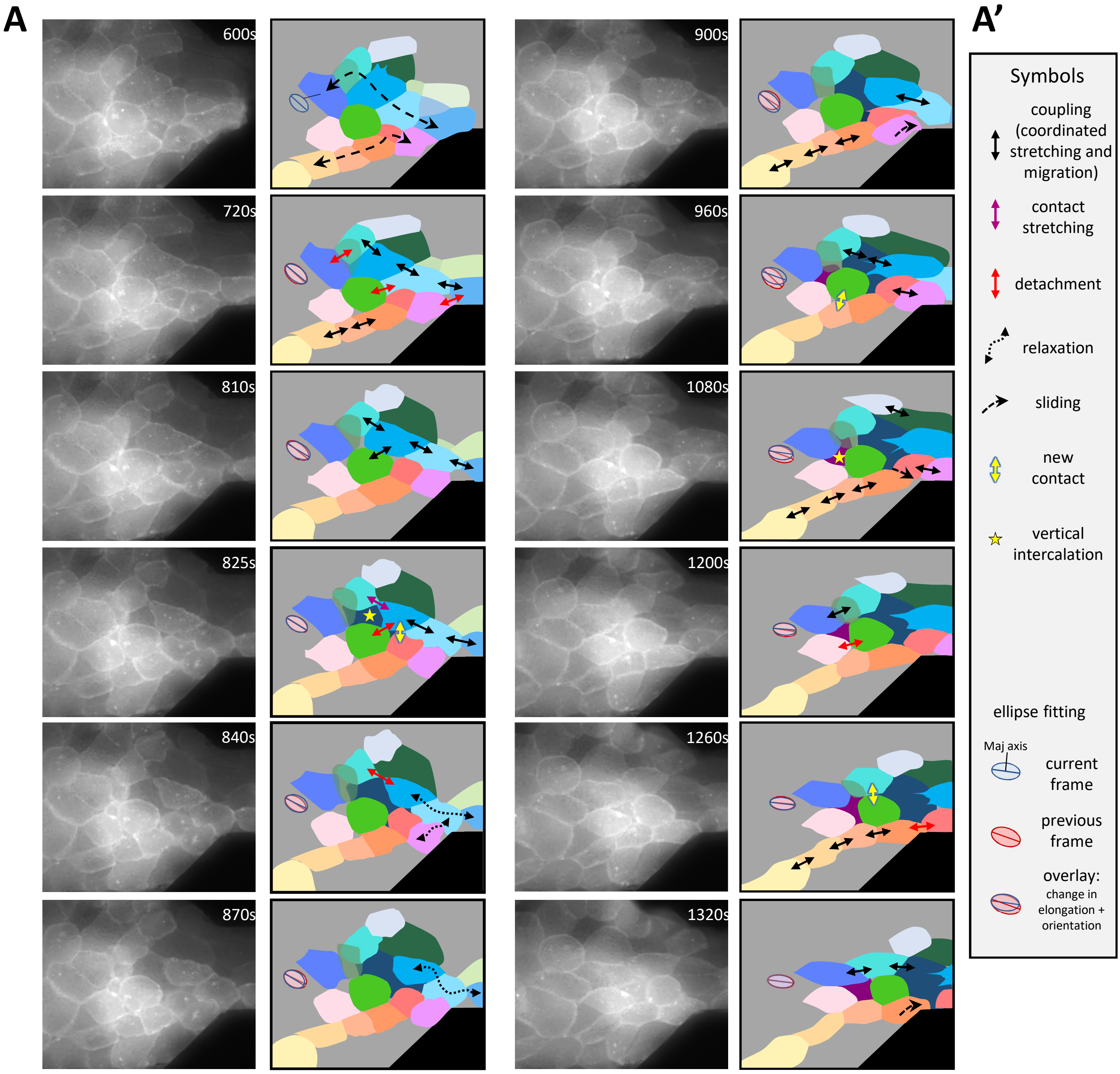
A
A'
600s
720s
810s
825s
840s
870s
900s
960s
1080s
1200s
1260s
1320s
Symbols
coupling (coordinated stretching and migration)
contact stretching
detachment
relaxation
sliding
new contact
vertical intercalation
ellipse fitting
Maj axis
current frame
previous frame
overlay: change in elongation + orientation

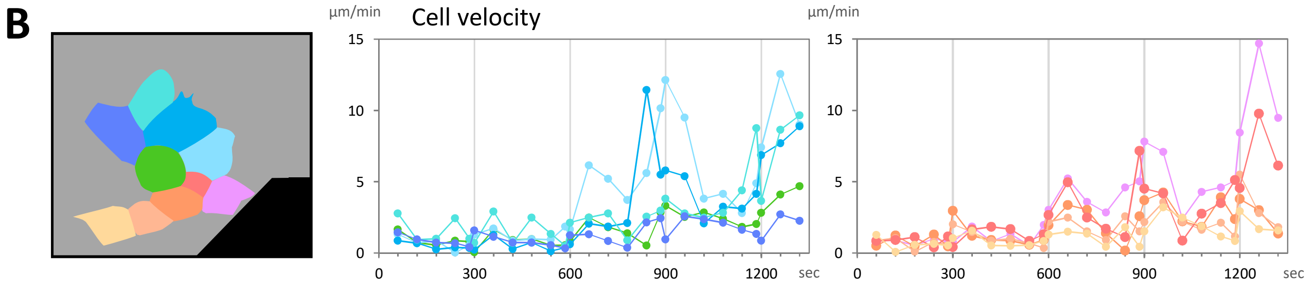
B
Cell velocity
µm/min
15
10
5
0
0
300
600
900
1200
sec

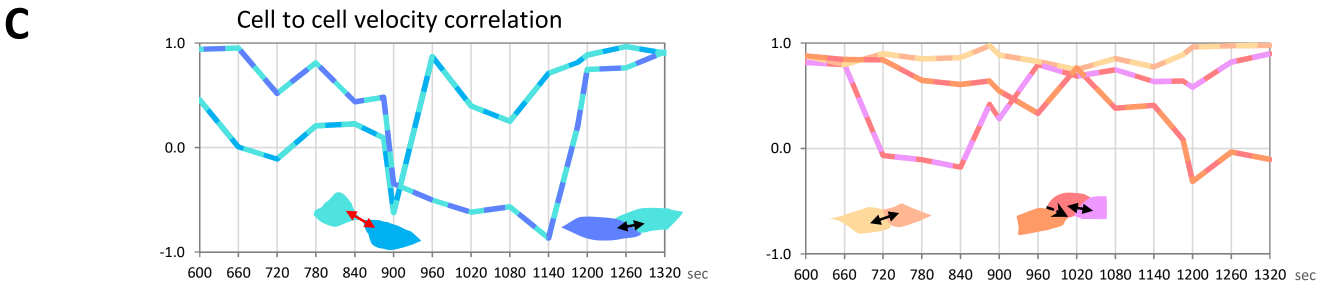
C
Cell to cell velocity correlation
1.0
0.0
-1.0
600
660
720
780
840
900
960
1020
1080
1140
1200
1260
1320
sec

**Supplemental Figure S5. Example of analysis of cell dynamics.**

**A)** Selected frames from tissue #11. A group cells positioned near the entrance of the channel was segmented thanks to the GFP membrane signal and tracked over time. Segmented cells are schematized in the right panels. Each cell is color-coded. The schemes are annotated with symbols pointing to stereotypical events such as cell-cell coupling (as indicated by coordinated stretching and migration) or relaxation, contact formation, stretching, sliding, and detachment, as indicated in A'. The two dashed double arrows in the first frame (600s) highlight two cell subgroups, respectively coded with shades of blue and shades of orange-red-pink that showed clear signs of cell-cell coupling along the axis of aspiration, and for which velocity measurements and correlations are presented in panels B and C. The schemes also include an ellipse corresponding to the adjacent purple-blue cell, fitted for the segmented cell shape as in Figures 2 and 3, here scaled down for clarity purpose. At each frame, a blue ellipse, fitted to the current frame, is superimposed on a pink ellipse corresponding to the previous frame. The line indicates the major axis. **B)** Cell velocities over the whole time-lapse, for the cells represented in the scheme on the left. **C)** Examples of cell to cell pairwise correlation of velocity during the sequence presented in A. For the blue row, a strong drop of correlation coincided with a transition from tight coupling to relaxation, which resulted from one contact detachment (840s, red double arrow). Coupling was recovered, apparently through the contribution of the dark blue cell that has started to appear through vertical intercalation in frame 823s (star). Eventually, the purple cell in the rear also became coupled (1200s) as shown both by the high raise of correlation (C) and by its reorientation and elongation (A, overlapping ellipses). As for the orange-red row, the rear cells remained strongly couple throughout the sequence (light oranges). In the front, remodelling involved sliding then detachment of the dark orange/red pair (coinciding with drop in velocity correlation after 1020s), while the red/pink pair underwent increasing coupling (see correlation in C, continuing on the channel outside of field in A).

### Figure S6

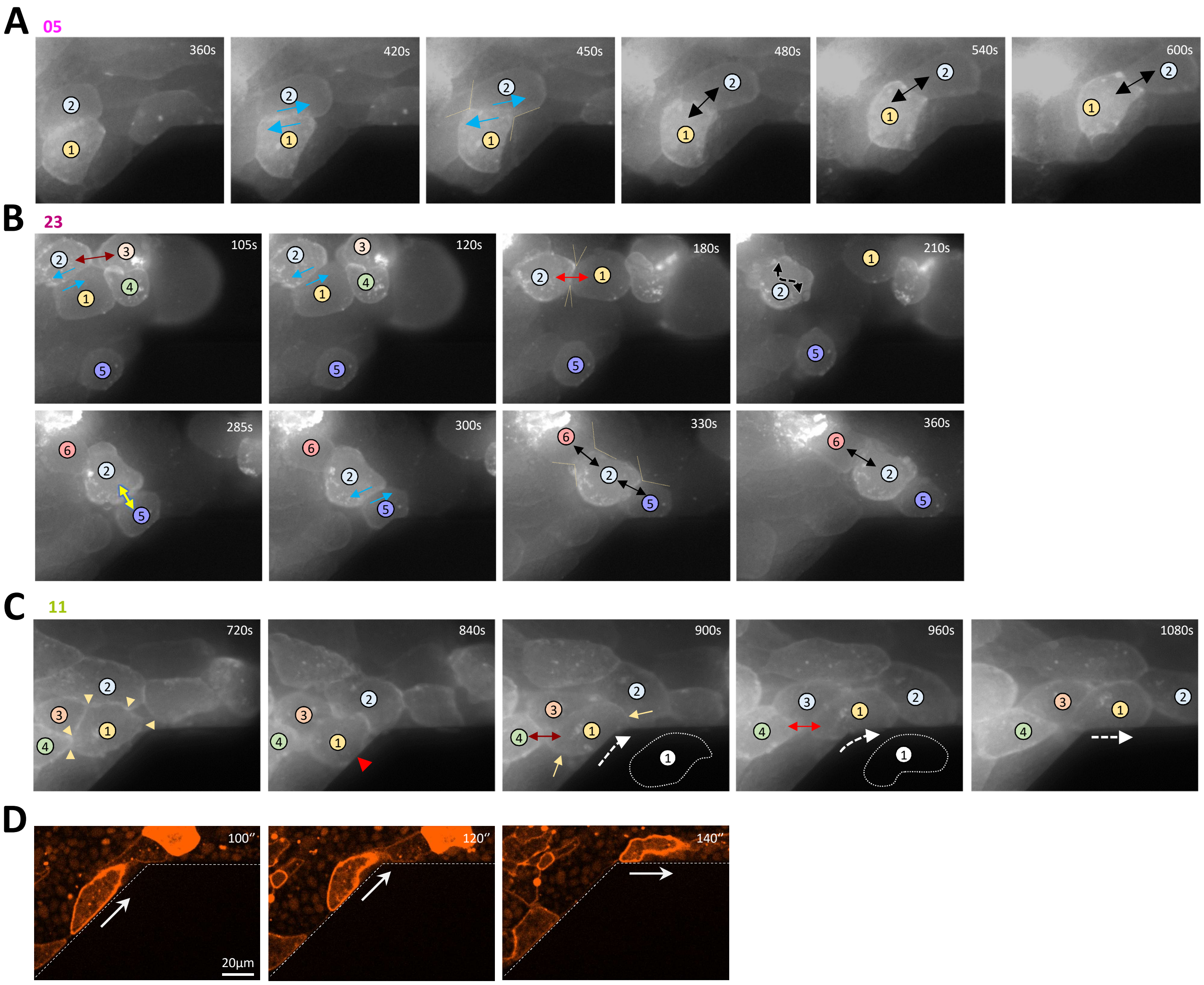


**Figure S6. Illustrations of typical cell movement behaviours.**
**A) Sliding and coupling.** Detail of tissue #05. 360-450s: Cell 1 and 2 first slide relative to each other (blue arrows). 450-480s: Transition from sliding to pulling, correlating with broad contact angles (yellow dotted lines), consisting with a "clamping" event. 480s onwards: both cells move together. They have elongated and aligned toward the channel axis. **B) Combination of various behaviours.** Detail of tissue #23. 105-120s: Cell 1 slides relative to cell 2 (blue arrows). Contact between cells 2 and 3 is stretched (dark red arrowhead) before detachment. 180s: Cell 1 has moved in front of cell 2. Acute contact angles are consistent with fast detachment ("peeling", see box 2). 210s: Contact 1-2 has detached. Cell 2 crumbled shape (symbolized by sinuous dashed black double arrow) is consistent with relaxation after detachment. 285s: Cells 2 and 5 establish contact (yellow double arrow), then slide relative to each other (300s, blue arrows). 330s: Cell 5 in front of cell 2. Coupling 2-5 and 2-6, with broad contact angles consistent with transient resistance, consistent with stretching of cell 6 at 360s. **C) Sliding slug.** Detail of tissue #11, related to figure 6. 720s: Cell 1 makes intimate contacts with neighbouring cells, displaying angular vertices (yellow arrowheads). 840-900s: Cell 1 gets in contact with the chamber wall (red arrowhead), its shape becomes rounder (yellow arrows, outlines reproduced as dotted white line on the right bottom), while it moves around the corner of the channel (dotted arrow). Contact with cell 4 is stretched (dark red and red double arrows) and breaks after 960s. **D)** Spinning confocal images of a sliding slug-like cell, expressing membrane Cherry.

### Figure S7

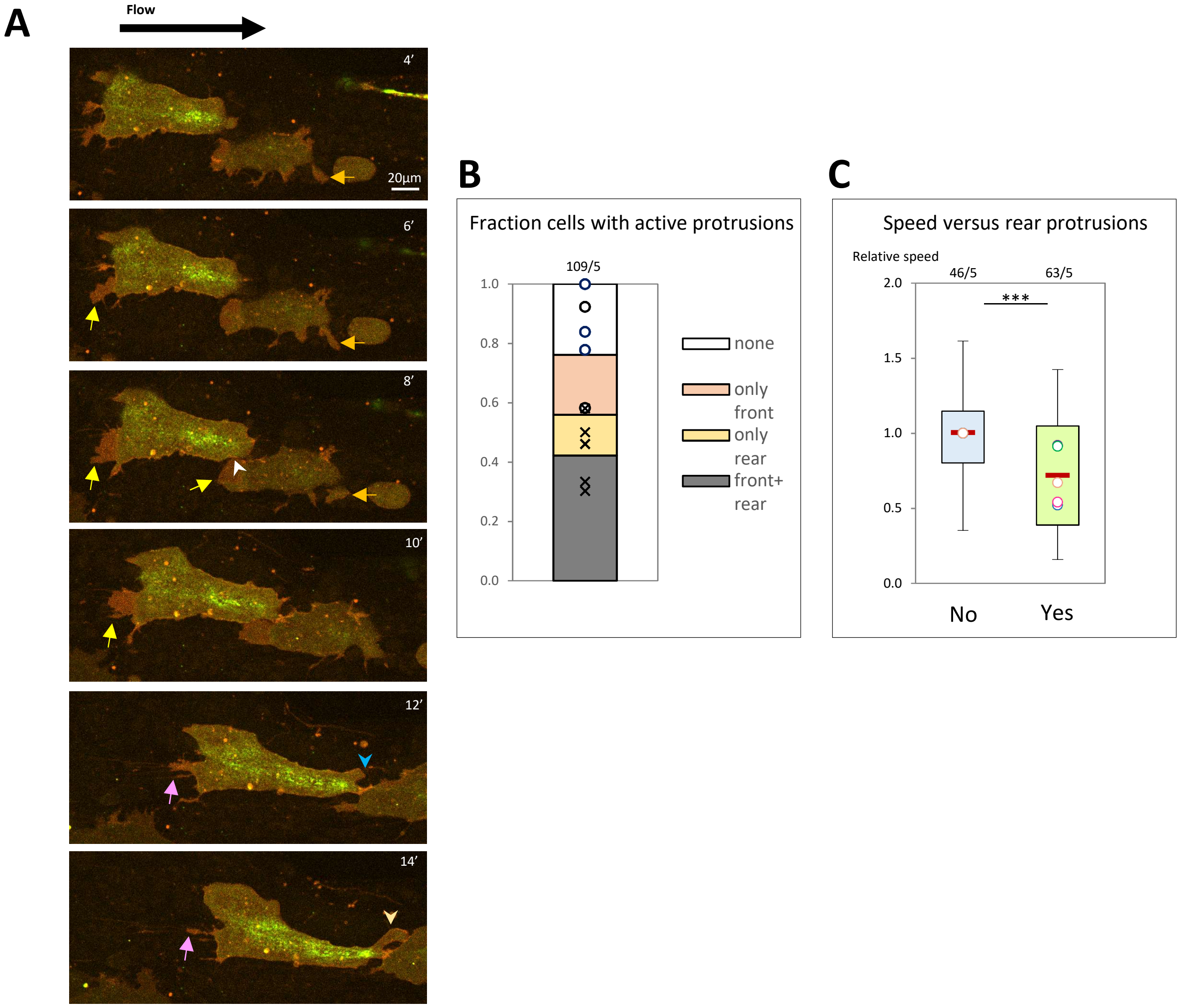


**Supplemental Figure S7. Analysis of protrusive activity.**

**A)** Spinning disk confocal imaging of cells expressing membrane-Cherry, co-expressed with myosin IIA-StayGold). Planes near the passivated glass are shown. Fields were chosen where expression was strongly mosaic, with single fluorescent cells highlighted over a background of unlabelled cells. Cells displayed multiple protrusions spreading, mainly directed toward the front and the back, that spread over neighbouring cells. Orange and yellow arrows point to protrusions that appear actively remodelled over time, respectively forwards or backwards. Pink arrows point to thin protrusions corresponding to retraction fibers, which appear as remnants of previously active protrusions. White arrowhead marks a new cell contact. Blue and yellow arrowheads point to a forming thick protrusion that establishes contact with the cell ahead. **B)** Quantification of the proportion of cells with protrusions at the front, at the rear, or at both ends. Only active protrusions were counted, thus excluding retraction fibers. Numbers on top: Total number of cells and number of experiments. Circles and crosses: Medians for each experiment, for the categories “no protrusions” and “both rear + front”. **C)** Cells with active rear protrusions display a significantly lower speed than the other cells, independent of the presence of front protrusions. Box plots correspond to the $1^{st}$ and $3^{rd}$ quartile, with the median. Whiskers, maximal and minimal values excluding outliers. Numbers on top: Total number of cells and number of experiments. Circles: Medians of individual experiments. Statistical analysis, two sided Student’s t-test. *** = p<0.001.
